# The Glycan-Specificity of the Pineapple Lectin AcmJRL and its Carbohydrate-Dependent Binding of the SARS-CoV-2 Spike Protein

**DOI:** 10.1101/2022.05.27.493400

**Authors:** Joscha Meiers, Jan Dastbaz, Sebastian Adam, Sari Rasheed, Susanne H. Kirsch, Peter Meiser, Peter Gross, Rolf Müller, Alexander Titz

## Abstract

The current SARS-CoV-2 pandemic has become one of the most challenging global health threats, with over 530 million reported infections by May 2022. In addition to vaccines, research and development have also been directed towards novel drugs. Since the highly glycosylated spike protein of SARS-CoV-2 is essential for infection, it constitutes a prime target for antiviral agents. The pineapple-derived jacalin-related lectin (AcmJRL) is present in the medication bromelain in significant quantities and has previously been described to bind mannosides. Here, we elucidated its ligand specificity by glycan array analysis, quantified the interaction with carbohydrates and validated high-mannose glycans as preferred ligands. Because the SARS-CoV-2 spike protein was previously reported to carry a high proportion of high-mannose N-glycans, we tested the binding of AcmJRL to recombinantly produced spike protein. We could demonstrate that AcmJRL binds the spike protein with a low micromolar K_D_ in a carbohydrate-dependent fashion, suggesting its use as a potential SARS-CoV-2 neutralising agent.

## Introduction

Since the end of 2019, the world is facing the severe acute respiratory syndrome coronavirus 2 (SARS-CoV-2) pandemic. SARS-CoV-2 is a novel coronavirus that rapidly spread all over the world and infected over 530 million people so far (May 2022). It can infect the respiratory tract and potentially results in the coronavirus-associated disease COVID-19. Especially for older or immunocompromised patients, COVID-19 is likely to be lethal. So far, more than 6.3 million deaths were reported in association with SARS-CoV-2.

A variety of novel and very potent vaccines entered the market at the end of 2020. Vaccination is an indispensable approach to protect society from a SARS-CoV-2 infection. However, a rising fraction of vaccinated individuals suffers from severe syndromes after infection due to a continuous adaptation of the virus. Furthermore, a very small fraction of people can not be vaccinated due to medical preconditions (1). Currently, first antiviral drugs like Remdesivir and Ritonavir/Nirmatrelvir (Paxlovid) are getting established in SARS-CoV-2 therapy and more are under investigation since we need novel pharmaceutical therapies to cure or prevent infections with SARS-CoV-2.

Coronavirus spike proteins (S-proteins) are essential for the infection process, they are trimerizing fusion proteins that consist of the two subunits S1 and S2. It has been shown that S-proteins have a complex and extensive glycosylation pattern (2), and coronavirus spike proteins typically contain between 23-38 *N*-glycosylation sites (3) per protomer with a significant population of oligomannose-type glycans (30%) (2). The elucidation of the glycosylation pattern (3) of the SARS-CoV-2 S-protein has been essential for the development of an effective vaccine (4). A site-specific glycan analysis by mass spectrometry has revealed that the 22 glycosylation sites on the S-protein monomer are occupied with a mixture of oligomannose-type, hybrid-type and complex-type glycans. The content of exclusively oligomannose-type glycosylation sites was determined to be 28%, which is above the level of typical host glycoproteins (3). However, it is less than for other viral glycoproteins like HIV-1 Env, where the amount of oligomannose-type glycans was found to be around 60% (3). *N*-glycosylation is not only vital for protein folding during protein expression, but these glycans also shield antigenic epitopes of S-protein and allow the virus to evade the host’s immune system (5). Further, glycans are important ligands for the first interaction with the host via its cell surface attachment receptors (5). For viral entry in airway epithelial cells, the receptor-binding domain (RBD) of SARS-CoV-2 S-protein binds angiotensin-converting-enzyme-2 (ACE2) with high affinity (6).

Given the extensive glycosylation of the S-proteins of coronaviruses, it has been hypothezised that targeting surface glycans of S-proteins could lead to a decreased virulence of SARS-CoV-2. In fact, a study revolving around 33 different plant carbohydrate-binding proteins, i.e. lectins, observed antiviral activity for 15 plant lectins against SARS-CoV and feline infectious peritonitis virus (FIPV) (7). Interestingly, the most prominent antiviral properties were found for mannose-specific lectins, which might be related to oligomannose-type glycans being essential for S-protein function. Further, Hoffmann et al. showed the ability of mammalian lectins (e.g. CLEC4G, CD209) to block SARS-CoV-2 infection *in vitro* (8).

Drug repurposing is a particularly interesting approach due to the acute nature of the current pandemic and led to the use of Remdesivir. Bromelain, a pineapple (*Ananas comosus*) stem extract, is an approved drug that shows anti-edematous, anti-inflammatory and fibrinolytic properties and is thus used to cure trauma-induced swelling (9), (10). Proteases, peptidic protease inhibitors and the mannose-binding lectin AcmJRL (also called AnLec) are the three main protein components of bromelain (11). It is likely that proteases are responsible for the anti-inflammatory properties of bromelain. Additionally, protease inhibitors prevent unspecific proteolysis as a safety mechanism that is slowly removed during the intake of bromelain. A putative mode of action of AcmJRL remains to be uncovered.

AcmJRL was recently characterised by Azarkan *et al*. (12) and its surprisingly high content in bromelain was determined by Gross *et al*. (11). AcmJRL belongs to the family of Jacalin-related lectins (JRL) (13). One of the first representatives of this family is jacalin, the lectin isolated from jackfruit (*Artocarpus integrifolia*). The JRL family can be divided in two main classes according to their ligand specificity (13). Galactose-specific JRL (gJRL) are found almost exclusively in the *Moraceae* plant family, most typically in the seed. Structurally, those JRLs are tetramers of four identical protomers, each containing one carbohydrate binding site. The complex biosynthesis of mature gJRLs includes co- and post-translational modifications from one preprotein including *N*-glycosylation. On the other hand, mannose-specific jacalin-related lectins (mJRL) are found in various plants. The structure of mannose-specific JRLs is less complex as they usually consist of two, four or eight unprocessed peptides. Due to the absence of a signal peptide, they are considered as cytoplasmic proteins.

Isothermal titration calorimetry experiments with AcmJRL revealed rather low binding affinity towards D-mannose (K_a_ = 178 M^−1^), D-glucose (K_a_ = 83 M^−1^) and Glc*N*Ac (K_a_ = 88 M^− 1^) (12). On the other hand, oligomannose structures like mannotriose (Man-α-1,6(Man-α-1,3)Man) and mannopentaose (Man-α-1,6(Man-α-1,3)Man-α-1,6(Man-α-1,3)Man) showed significantly higher binding affinities of K_a_ = 734 M^−1^ to 1694 M^−1^.

Like other mannose-specific JRLs, AcmJRL adopts a characteristic β-prism fold and two monomers align side-by-side forming a dimer. Although a tetrameric form of AcmJRL could also be assigned from the monomers in the asymmetric unit, this is likely a crystallization artefact (see below). In co-crystal structures with D-mannose and methyl α-D-mannopyranoside, two carbohydrates are bound by one monomer in a conserved binding pose. Overall, the interactions are comparable to the binding of D-mannose by BanLec, a closely related mannose-specific JRL from banana (*Musa acuminata*). BanLec is reported as a potent viral entry inhibitor of HIV-1, HCV and influenza virus (14,15). However, the mitogenic activity of native BanLec limits its therapeutic use. Interestingly, the structure of AcmJRL shares similarities with a genetically engineered BanLec (15) with reduced mitogenic activity. Thus, Azarkan *et al*. postulated a potential use of AcmJRL as an alternative to BanLec against mannosylated viruses.

Here, the ligand specificity of AcmJRL was further characterised with two glycan arrays and the interaction was quantified in a competitive binding assay. In a next step, we demonstrated the ability of AcmJRL to bind to the SARS-CoV-2 spike protein as well as its isolated receptor-binding domain (RBD) with low micromolar affinity in a carbohydrate-dependent manner. Further, we showed that the binding of the spike RBD to its receptor ACE2 can be inhibited by AcmJRL. Consequently, it is possible that the mannose-binding lectin AcmJRL can neutralise the SARS-CoV-2 virus through binding to its spike protein.

## Results and discussion

### Isolation and characterisation of AcmJRL from bromelain

The mannophilic lectin AcmJRL was isolated by affinity chromatography from pineapple stem extract (bromelain) on a mannosylated stationary phase according to the procedure reported by Azarkan *et al*. (12). Prior to purification, the soluble protein fraction of bromelain was obtained by aqueous extraction in presence of S-methyl methanethiosulfonate to block bromelain’s high proteolytic activity. Using mannosylated sepharose beads (16) 0.9 - 1.6 mg AcmJRL were obtained per gram bromelain after elution with mannose.

The identity of the isolated protein was confirmed by SDS-PAGE (Figure 1A) and mass spectrometry (average molecular mass = 15346 Da, Figure 1B). The main peak (*m/z* = 15388) can be assigned to an acetonitrile adduct [M+H+MeCN]^+^ of the AcmJRL monomer. As reported by Gross *et al*. (11), two additional mass peaks were observed in a ratio of 100 : 65 : 17, separated by a mass shift of 162 Da. During the industrial production of bromelain, the pineapple stem extract is loaded on maltodextrin, a hydrolysis product of starch, to simplify its handling. Thus, it contains carbohydrates like glucose and glucooligo-saccharides that can react with primary amines of proteins to form Schiff-bases, which is presumably followed by an irreversible Amadori rearrangement (Figure S1) towards a stable α-amino ketone corresponding to advanced glycation end-products (17). This glycation results in a mass increase of 162 Da that was observed in the MS-spectrum (Figure 1B). The presence of two signals of +162 Da and +324 Da suggests that this reaction occurred twice on the protein or one disaccharide of maltose reacted. However, it is not clear if it is a statistical mixture or if two specific lysines were affected by this reaction (5 lysines are present in AcmJRL). It was not specified whether the AcmJRL isolated and crystallised by Azarkan *et al*. was also glycated. Inspection of the electron density map of crystallised AcmJRL (PDB: 6FLY (12)) did not show evidence for unassigned electron density which could also result from a high flexibility at the protein surface or a statistical distribution of the glycation.

**Figure 1.**
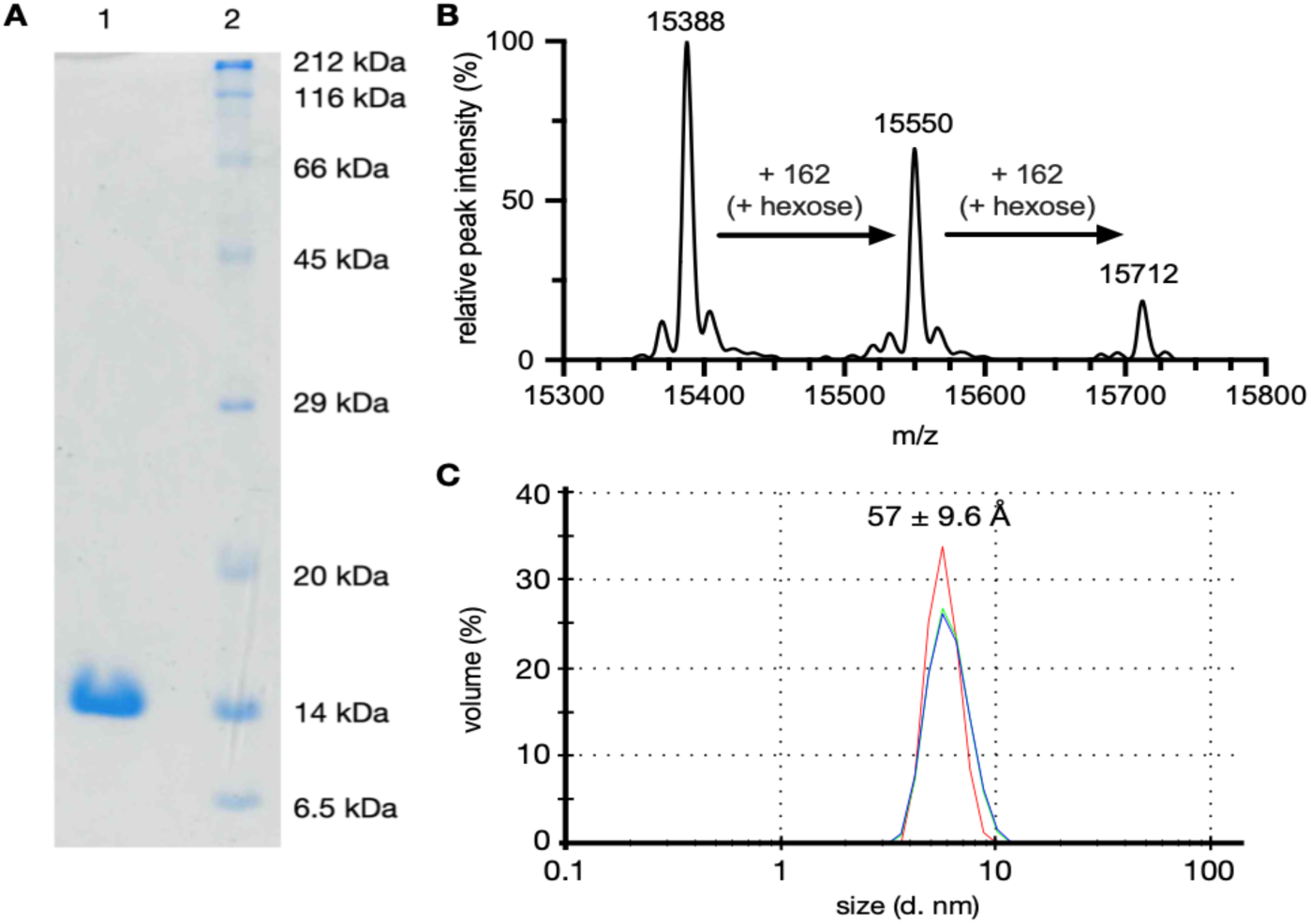
**A** Affinity purified AcmJRL (lane 1) and molecular weight marker (lane 2) analysed by SDS-PAGE (18%). **B** ESI-MS spectrum of AcmJRL after maximum entropy deconvolution. Main peak (m/z = 15388) corresponds to an acetonitrile adduct [M+H+MeCN]^+^. Peaks at *m/z* = 15550 and *m/z* = 15712 most likely result from glycation (Figure S1). **C** Dynamic light scattering analysis of AcmJRL (size distribution by volume). Peak at 57 ± 9.6 Å indicates a dimerisation in solution.

Previous studies showed dimerisation of AcmJRL in solution, determined by size exclusion chromatography and equilibrium unfolding experiments (12). However, Azarkan *et al*. showed that AcmJRL crystallises as a tetramer (see introduction). We therefore used dynamic light scattering (DLS) to determine the hydrodynamic diameter of AcmJRL in buffered solution (Figure 1C). The measured hydrodynamic diameter of 57 ± 9.6 Å corresponds to the radius of the dimer (Figure S2), rather than to monomers or tetramers. Additionally, our differential scanning fluorimetry studies suggested two unfolding events at T_1_ = 58 - 60°C and T_2_ = 73 - 74°C (Figure S2) which could reflect dissociation of the dimer followed by protein denaturation.

### Glycan array analysis of AcmJRL

In general, jacalin-related lectins can be clustered by their carbohydrate specificity into galactophilic and manno-/glucophilic subgroups. AcmJRL is reported to have a millimolar affinity towards mannosides and glucosides (12). Glycan microarrays (18) allow the simultaneous binding specificity analysis of carbohydrate-binding proteins on a library of oligosaccharides, using e.g. fluorescence-labelling for optical quantification.

Residues Lys50 and Lys63 are most accessible in AcmJRL for labeling with amine reactive dyes according to the solvent accessible protein surface calculated from the crystal structure (Figure S4). Fluorescence labelling with FITC resulted in a rather low labelling efficiency of 0.44. Since low labelling efficiency combined with high photobleaching of fluorescein could result in poor sensitivity, we shifted to the Cyanine-3 (Cy3) dye. The latter is a bright, photostable and pH-insensitive orange fluorescent dye. Unfortunately, Cy3-labelling via its NHS active ester also resulted in a low labelling efficiency of 0.55. Notably, one or two of the five lysines of AcmJRL are not available for labelling as they are blocked by glycation (Figure 1B). Further, the reduced accessibility of Lys6, Lys29 and Lys79 could explain the low labelling yields. On the other hand, only ca. 10% of AcmJRL is bis-glycosylated, 36% is mono-glycosylated and 54% is unmodified (Figure 1B).

To elucidate its glycan specificity on a larger collection of oligosaccharides, AcmJRL was analysed on the Consortium for Functional Glycomics mammalian glycan array (19) in an updated version with 585 distinct carbohydrate epitopes (Figure 2A, Tables S5 and S6). In agreement with its known mannose specificity, the lectin showed a high affinity towards several α-mannosides (e.g. CFG-GLYCAN ID **310**, Man-α-1,6(Man-α-1,3)Man-α-1,6(Man-α-1,3)Man-β, RFU = 410 ± 141).

**Figure 2.**
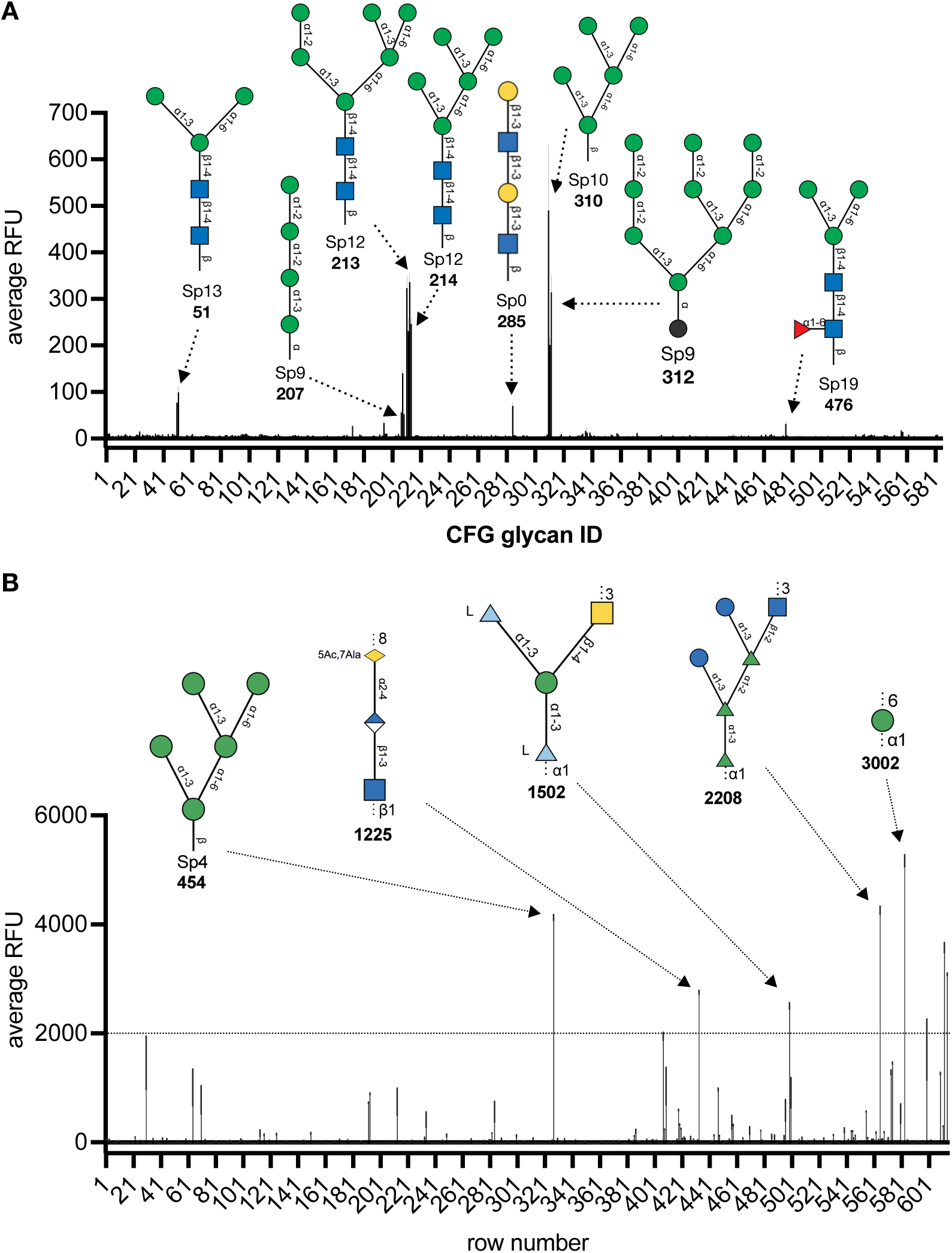
Glycan array analysis of AcmJRL: **A** CFG mammalian glycan array tested with FITC-labeled AcmJRL at 50 μg/mL, **B** Semiotic glycan array tested with Cy3-labeled AcmJRL at 20 μg/mL. Data is presented as average RFU ± s.d. from 6 replicates on array. RFU intensities differ between arrays since different fluorophores, protocols and scanners were used. **Number** = glycan ID. Translation from row number to Semiotik glycan ID and short name can be found in tables S1 and S2.

Bi- and triantennary mannosides showed higher apparent binding affinities towards AcmJRL than monovalent glycans (e.g. compare CFG-GLYCAN ID **312** vs. **207**/**209**). As described above, these oligomannosidic structures can be found in N-glycans of mammalian proteins and have been reported for viral surface proteins such as the SARS-CoV-2 spike protein. Increased binding by multivalent ligand presentation is common for glycan-lectin recognition. The mannotriose epitope Man-α-1,6(Man-α-1,3)Man (commonly present in CFG-GLYCAN ID **211, 51** and **50**), the mannopentaose epitope Man-α-1,6(Man-α-1,3)Man-α-1,6(Man-α-1,3)Man (commonly present in in CFG-GLYCAN ID **310** and **214**) and the mannohexaose epitope Man-α-1,6(Man-α-1,3)Man-α-1,6(Man-α-1,2-Man-α-1,3)Man (present in GFG-GID **213**) showed highest binding responses which corresponded to reported data (K_d_ mannotriose = 1.4 mM, K_d_ mannopentaose = 590 μM) (12). One single glycan hit without mannose but a terminal β-galactoside in a LacNAc repeat (CFG-GLYCAN ID **285**, Figure 2A) is presumably a false positive due to dose-independent changes in signal intensity (5 vs 50 μg/mL).

AcmJRL forms a dimer in solution and can bind up to two mannosides per binding site. The distance from the binding site of one monomer to the other monomer in the crystal structure is approx. 46-50 Å. The distance between the anomeric carbons of two mannosides within one binding site is approx. 14 Å (from C1 to C1), which is similar to the distance between two mannoses in mannopentaose suggesting a possible chelation binding mode by this ligand. On the other hand, mannopentaose could also preorganise two α-mannosides in a way that allows the rapid rebinding with the two binding sites within one monomer.

Monovalent α-glucosides showed very low, but significant binding (e.g. CFG-GLYCAN ID **195**, Glc-α-1,4-Glc-β, RFU = 33 ± 5). Unfortunately, no multivalent glucosides are available on this array to understand the influence of multivalency for these epitopes.

In addition to the CFG glycan array, we analysed AcmJRL on the Semiotik glycan array(20), which also features mammalian glycans and additionally a large variety of other glycans, mainly from bacterial species (Figure 2B). Although mannopentaose (Semiotik glycan ID (SGID) **454**) and poly-Man-α-1,6 (mannan, SGID **3002**) could be confirmed as a ligand for AcmJRL, the smaller mannotriose (SGID **258**) showed no binding, arguably due to a shorter linker length (Sp4) preventing accessibility for the protein. In contrast to the CFG glycan array, a multivalent α-glucoside is present on the Semiotik chip (SGID **2208**) and was well recognized by the lectin, underlining the affinity of AcmJRL towards α-glucosides. Interestingly, two unrelated bacterial O-antigens were also recognised: *E. coli* O161 (SGID **1225**, -8(D-Ala1,7)Leg5Ac-α-2,4-GlcA-β-1,3-GlcNAc-β1-) and *A. hydrophila* O34deAc (SGID **1502**, -3GalNAc-β-1,4(L-6dTal-α-1,3)Man-α-1,3L-6dTal-α-1-). However, SGID **1502** showed a nonlinear dose-response, which requires orthogonal binding assays for verification.

### Development of a competitive binding assay for AcmJRL

Glycan arrays provide valuable insight into carbohydrate specificity in high-throughput. For a quantitative binding analysis, we developed a solution phase competitive binding assay based on our previous work for other lectins using fluorescence polarisation (21, 22, 23). Fluorescein-labelled α-D-mannoside **1** (24) was titrated with AcmJRL and a K_d_ of 58.9 ± 5.9 μM was determined (Figure 3). This rather high affinity of AcmJRL for this α-mannoside was surprising, compared to the lectin’s reported affinity towards the previously best known ligand mannobiose Manα1-3Man (K_d_ = 2.4 mM (12)). Interestingly, such a discrepancy between the weaker carbohydrate alone and a boost in binding affinity for its fluorophore-labeled derivative was already observed for the bacterial lectin PllA (K_d_ = 62.7 μM and K_d_ 520 μM, for FITC-modified α-D-Gal and Me-α-D-Gal, respectively) (24).

**Figure 3.**
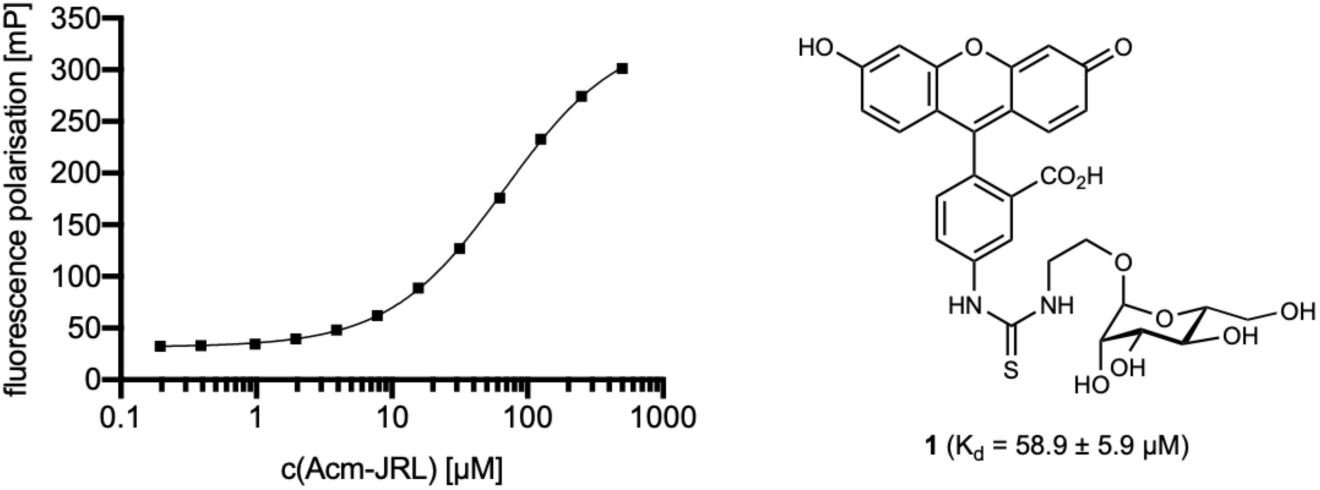
Direct binding of AcmJRL to mannose-based fluorescent ligand **1** and quantification using fluorescence polarisation. Dissociation constant and standard deviations were obtained from three independent experiments of triplicates on each plate.

This system was used to screen several carbohydrates in competitive binding assays. Next to the carbohydrate hits D-mannose and D-glucose and the two non-recognised epitopes D-galactose and L-fucose from the glycan arrays, other plant carbohydrates like L-rhamnose (Rha), D-xylose (Xyl) and D-arabinose (Ara) were tested (Figure 4A). First, single concentration inhibition assays again confirmed the affinity of AcmJRL towards α-mannosides and α-glucosides (Figure 4A). Additionally, glucosamine (GlcN) and *N*-acetyl-glucosamine (GlcNAc) were confirmed as ligands. In accordance with the glycan array results, none of the other tested carbohydrates inhibited AcmJRL at 10 mM. In order to evaluate the influence of the glycosidic linkage of the oligomannosides (α-1,2 *vs* α-1,3 *vs* α-1,6), a titration of the appropriate dimannosides was performed. Only subtle affinity differences between the isomers could be observed (Figure 4B, IC_50_ (Man-α-1,2-Man) = 9.1 ± 0.8 mM, IC_50_ (Man-α-1,6-Man) = 11.3 ± 1.7 mM), with Man-α-1,3-Man (IC_50_ = 6.9 ± 0.6 mM) having the highest potency. Comparing the glycan array results with this finding could explain why mannopentaose which contains two Man-α-1,3-Man-epitopes was preferentially recognised over other bis-(Man-α-1,2-Man)-presenting epitopes (e.g. CFG-GLYCAN ID **208**). Interestingly, Me-α-D-glucoside (IC_50_ = 6.8 ± 0.5 mM) showed higher affinity towards AcmJRL than Me-α-D-Man (IC_50_ = 10.8 ± 1.8 mM) and the di-mannosides (Figure 4B). Furthermore, from our single point inhibition experiments it is evident that the glycosidic linkage in glucosides is crucial for recognition by AcmJRL: maltobiose (Glc-α-1,4-Glc) showed highest inhibition, whereas cellobiose (Glc-β-1,4-Glc) had no effect.

**Figure 4.**
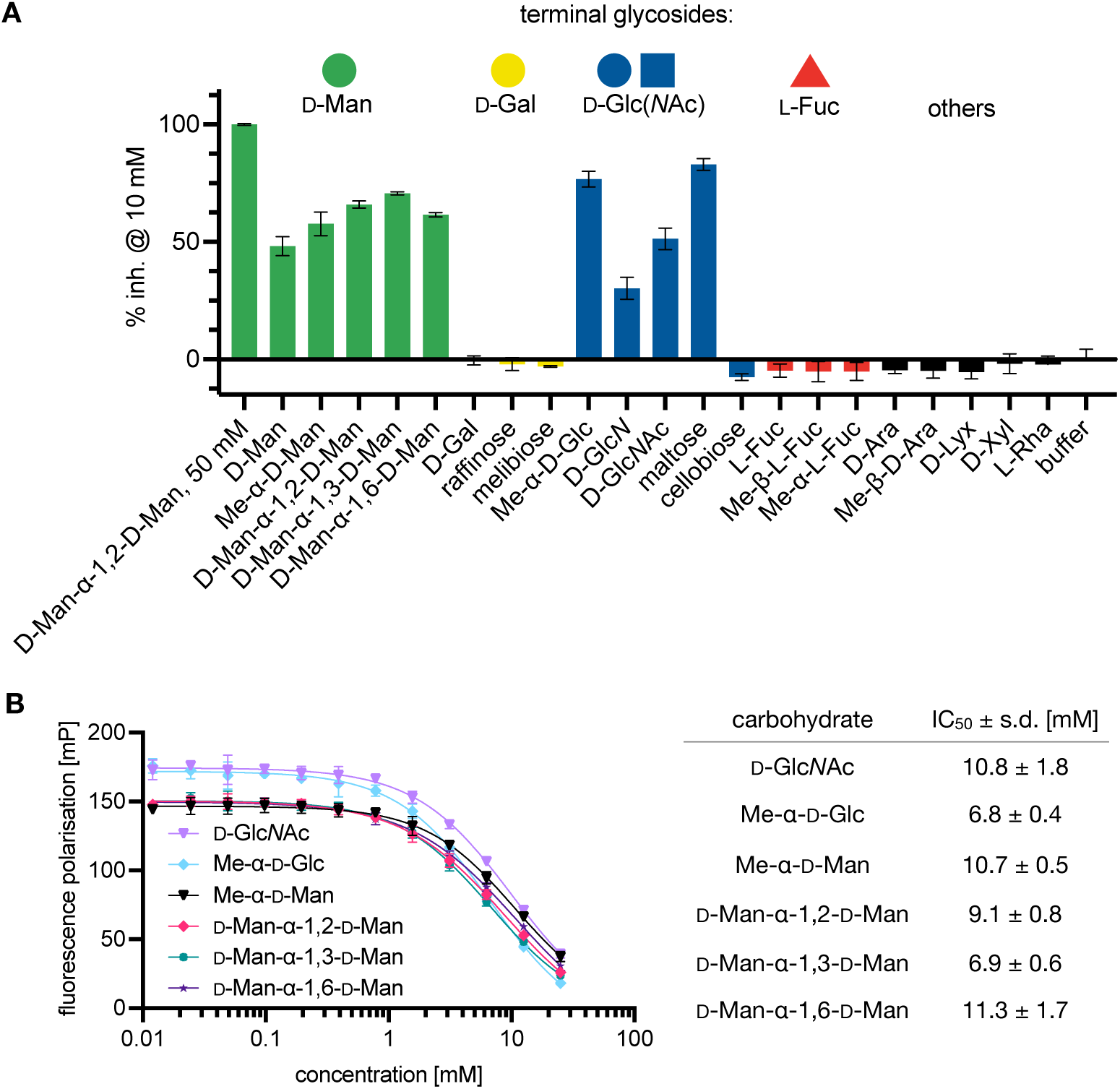
Competitive binding of various carbohydrate ligands to AcmJRL: **A** Inhibition of AcmJRL by a carbohydrate panel at 10 mM. Fluorescence polarisation in presence of 50 mM Man-α1,2-Man was defined as 100% inhibition. **B** Dose-response curves of AcmJRL with differently linked mannosides (left) or Me-α-Glc, Glc*N*Ac. Data is shown as mean ± s.d. from technical triplicates on plate.

### Expression of SARS-CoV-2 Spike Protein

Given the potential antiviral properties of plant lectins, our characterisation of AcmJRL made us speculate about the binding of AcmJRL to the heavily glycosylated SARS-CoV-2 spike protein.

The spike-protein of SARS-CoV-2 was recombinantly produced in HEK293 cells using a pCAGGS-based eukaryotic vector system yielding 1.42 mg glycoprotein from one batch of 560 mL. Identity and purity of the protein were determined by gel electrophoresis (Figure 5A) and mass spectroscopy (Figure 5B). Both experiments coherently show a molecular weight around 145 kDa. Interestingly, mass spectroscopy revealed four major masses after maximum entropy (MaxEnt) deconvolution of the centroided mass spectrum, each separated by approximately 2 kDa (Figure 5B). As described above, S-protein usually exhibits highly complex glycosylation, which leads to a non-homogeneous sample. Therefore, the heterogeneous mass distribution determined by ESI-MS presumably resulted from the presence of different glycoforms.

**Figure 5.**
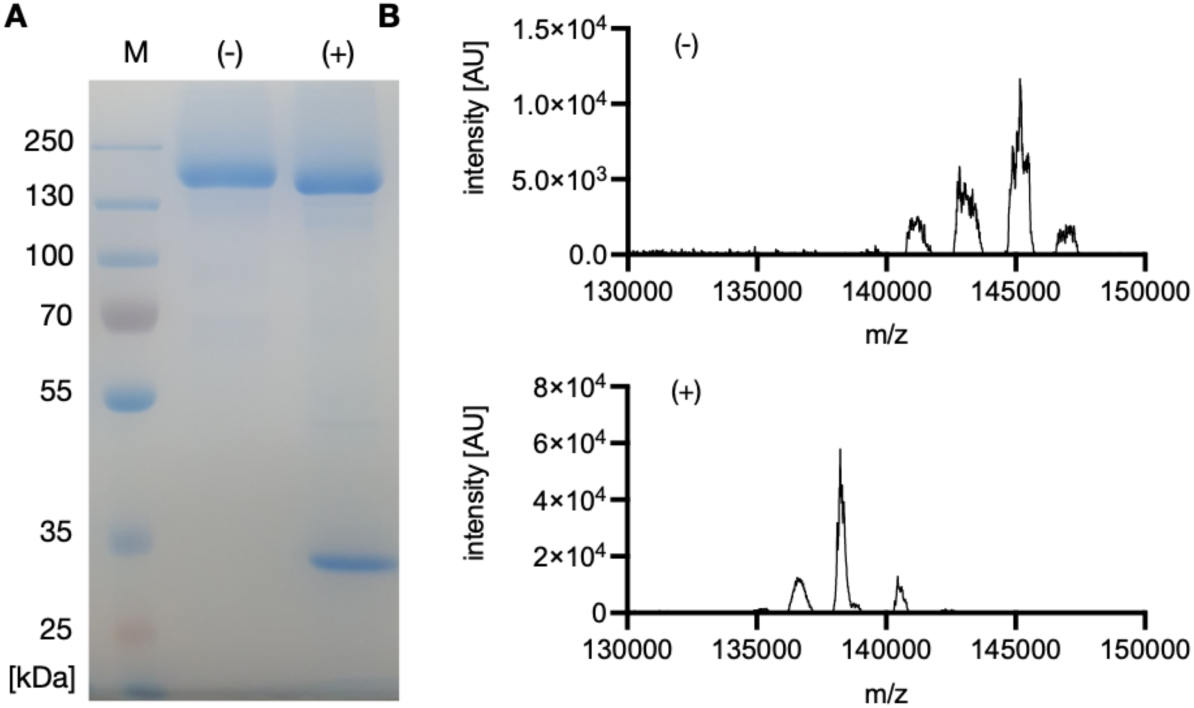
**A** SDS-PAGE (10%) of native S-protein (-) and S-protein digested with PNGase F (+). **B** Deconvoluted mass spectrum of native S-protein (-) and S-protein digested with PNGase F (+). In both experiments, a mass shift of several kDa is visible after treatment with PNGase F indicating removal of N-glycans.

Peptide-N-glycanase F (PNGase F) is an amidase that specifically hydrolyses amide bonds between the reducing end Glc*N*Ac and asparagine residues of *N*-glycans. Consequently, PNGase F was used to verify the glycosylation of the recombinantly produced *N*-glycosylated S-protein. S-protein was incubated in the presence of PNGase F and compared with the untreated protein sample (Figure 5). In fact, the treatment resulted in a faster migration on the SDS-gel indicating a molecular weight reduced by several kDa (Figure 5A). Coherently, the MaxEnt deconvoluted mass spectrum of the treated S-protein shows one major peak around 138 kDa, together with two small satellite peaks. Notably, all mass peaks of the PNGase F-treated species were much sharper and reached higher signal intensities compared to the native protein. The lower intensity of the deconvoluted mass peaks of the untreated protein in comparison to the higher intensity of the deconvoluted mass peaks of the PNGase F-treated protein indicated that a smaller diversity of glycoprotein species is existent in the deglycosylated sample while in the glycosylated native sample, the presence of a complex mixture of different glycoforms is responsible for the lower peak intensities due to the lower abundance of each individual species. The most intense peak in the deconvoluted mass spectra reveals a mass shift of approximately 7 kDa, resulting in a main mass peak around 138 kDa. Therefore, the mass spectrometric analysis of the enzyme treated sample qualitatively confirmed an extensive glycosylation of the recombinantly produced glycoprotein.

### Mannose-dependence of AcmJRL binding to SARS-CoV-2 spike-protein

After verification of the S-protein identity, a biophysical assay for determination of binding kinetics and affinity of AcmJRL to the S-protein was established. Given the high molecular weights of both interaction partners, Surface Plasmon Resonance (SPR) analysis was chosen to determine binding affinity and kinetics of AcmJRL against immobilised S-protein. Single cycle kinetics were performed by injecting AcmJRL at increasing concentrations from 2.5 to 20 μM onto a sensor chip with immobilised S-protein and a dose-dependent response was observed. Even at the highest concentration, saturation could not be observed after 120 sec association, hinting at an extensive number of available binding sites for AcmJRL. In addition, only an incomplete dissociation was recorded during the dissociation phases. The association rate (*k*_*on*_) was 1057 ± 94 M^−1^s^−1^, the dissociation rate (*k*_*off*_) was 3.42 ± 0.4 × 10^−3^ s^−1^ which corresponds to a K_D_ of 3.27 ± 0.5 μM. A similar K_D_ of 11.1 ± 2.5 μM was determined after fitting the response after 115 sec contact time of the association with the Langmuir isotherm, which is more error prone due to the fact that saturation was not reached.

Interestingly, the K_D_ increased gradually for each single experiment performed for the technical replicates. (Dissociation constants calculated from Langmuir isotherm: 7.9 μM, 11.5 μM, 14.0 μM; Dissociation constants calculated from rate constants: 2.7 μM, 3.1 μM, 4.0 μM). This observation likely resulted from some AcmJRL remaining bound during the regeneration cycles as the S-protein has a vast number of possible binding sites in its numerous N-glycans for AcmJRL. Furthermore, during dissociation phases the RU (response units) values did not fully decrease to the baseline response, indicative for an incomplete dissociation of AcmJRL.

As AcmJRL interacts with the mannotriose epitope (Man-α-1,6(Man-α-1,3)Man, present in CFG glycan ID **211, 213, 51** and **50**) in the glycan array analysis, single cycle kinetics of AcmJRL with the addition of 10 mM mannotriose in the sample buffer were also conducted to analyse the glycan-dependence of the AcmJRL-spike binding. The loss of observable binding to the spike protein in presence of the competitor demonstrates the mannose-dependent binding of AcmJRL (Figure 6).

**Figure 6.**
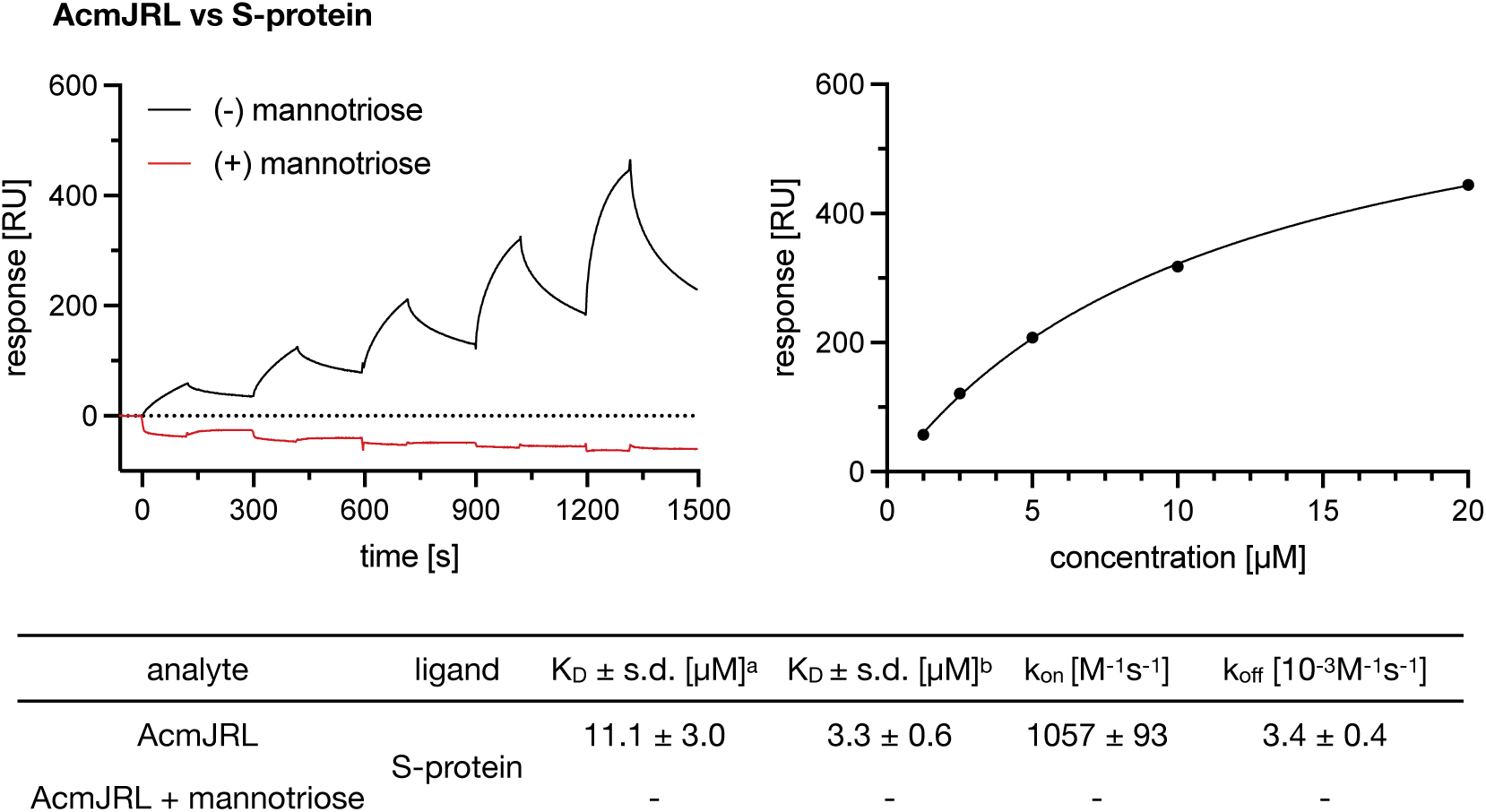
Surface plasmon resonance (SPR) sensorgrams of the binding of AcmJRL to immobilised S-protein in absence (black) and in presence of 10 mM mannotriose (Man-α-1,6(Man-α-1,3)Man, red). Langmuir graph of AcmJRL binding to S-protein. Dissociation constant calculated from ^a^Langmuir isotherm or ^b^rate constants, K_D_= *k_off_*/*k_on_*.

### AcmJRL binds to SARS-CoV-2 spike RBD and to its native receptor ACE2

The RBD of the S-protein is essential for the SARS-CoV-2 infection process, as it mediates binding to the human cell surface receptor ACE2, allowing the attachment of the virus. As we observed the binding of AcmJRL to the S-protein, we set out to determine if the lectin is able to block this essential mechanism for the infection through binding to the spike protein. Complex glycosylation is present in many other human cell surface proteins, such as the receptor ACE2.

Consequently, we also analysed the binding of AcmJRL to ACE2 as well as to spike RBD produced for interaction studies. Single cycle kinetics on SPR were performed for AcmJRL binding to both the S-protein RBD and the human ACE2 receptor (Figure 7). Twentyone glycosylation sites are present in full length S-protein (per monomer, 63 in the trimeric form) literally leading to a sweet fur covering the entire spike (25, 26), two of which reside within its RBD (5). The observed association of AcmJRL to S-protein RBD reflects this reduced glycosylation well, as saturation was now reached during association. The determined K_D_ value of 12.9 μM from kinetics or 35 μM from the Langmuir plot is about threefold higher compared to its interaction with full length S-protein. The association kinetics (*k*_*on*_ = 1490 ± 455 M^−1^s^−1^) were comparable to those for AcmJRL binding to S-protein (Figure 6). However, the dissociation of AcmJRL from spike RBD is about fourfold faster, displayed in the determined *k*_*off*_ = 19.4 ± 6.7 × 10^−3^ M^−1^s^−1^, a consequence of the reduced extent of glycosylation of the RBD.

**Figure 7.**
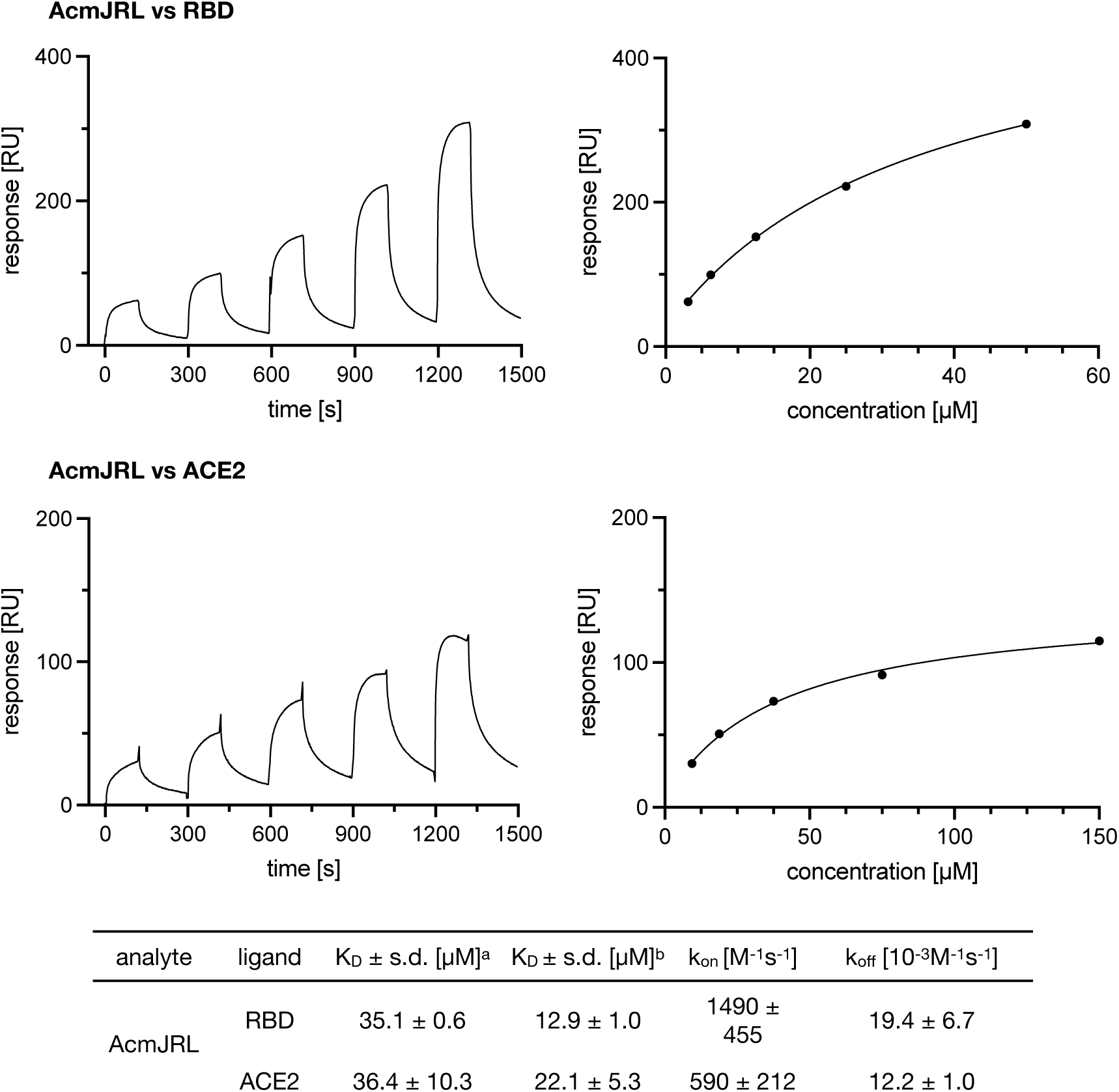
Surface plasmon resonance (SPR) sensorgrams with Langmuir graphs of the binding of AcmJRL to S-protein RBD (top) and ACE2 receptor (bottom). Dissociation constant calculated from ^a^Langmuir isotherm or ^b^rate constants, K_D_= *k_off_*/*k_on_*.

Human ACE2 is also highly glycosylated, carrying more matured complex glycans compared to the high-mannose enriched virus surface protein (27). From SPR analysis of AcmJRL binding to immobilised recombinant ACE2 produced in HEK293 cells, we obtained a K_D_ value of 22.1 ± 5.3 μM calculated from the Langmuir isotherm, which is sevenfold higher than the one obtained for binding to the S-protein. This observation could be a direct result of the altered glycosylation pattern of this human receptor that is distinct from the viral proteins. The association rate of AcmJRL to ACE2 (*k*_*on*_= 590 ± 212 M^−1^s^−1^) was also slower compared to the one for the S-protein. Further, the threefold higher dissociation rate (*k*_*off*_ 12.2 ± 1.0 × 10^−3^ M^−1^s^−1^) of AcmJRL from the ACE2 receptor indicates a faster dissociation of the complex. In contrast to the interaction with full length S-protein, complete dissociation of AcmJRL from both the RBD and ACE2 complexes was observed. The observed affinity of AcmJRL for both S-protein and ACE2 could therefore result in a synergistic inhibitory effect on viral cell entry.

### AcmJRL weakens the interaction of spike RBD with ACE2

The interaction between the spike RBD and its ACE2 receptor is characterised by nanomolar affinity and is essential for the infection process. As AcmJRL binds the S-protein RBD and ACE2 with low micromolar affinity, it is likely that the spike interaction with ACE2 could be inhibited by AcmJRL. To test this hypothesis, we first reproduced the affinity reported for the S-protein RBD to the immobilised ACE2 receptor (Figure 8) by SPR and obtained a K_D_ of 10.9 nM (28).

**Figure 8.**
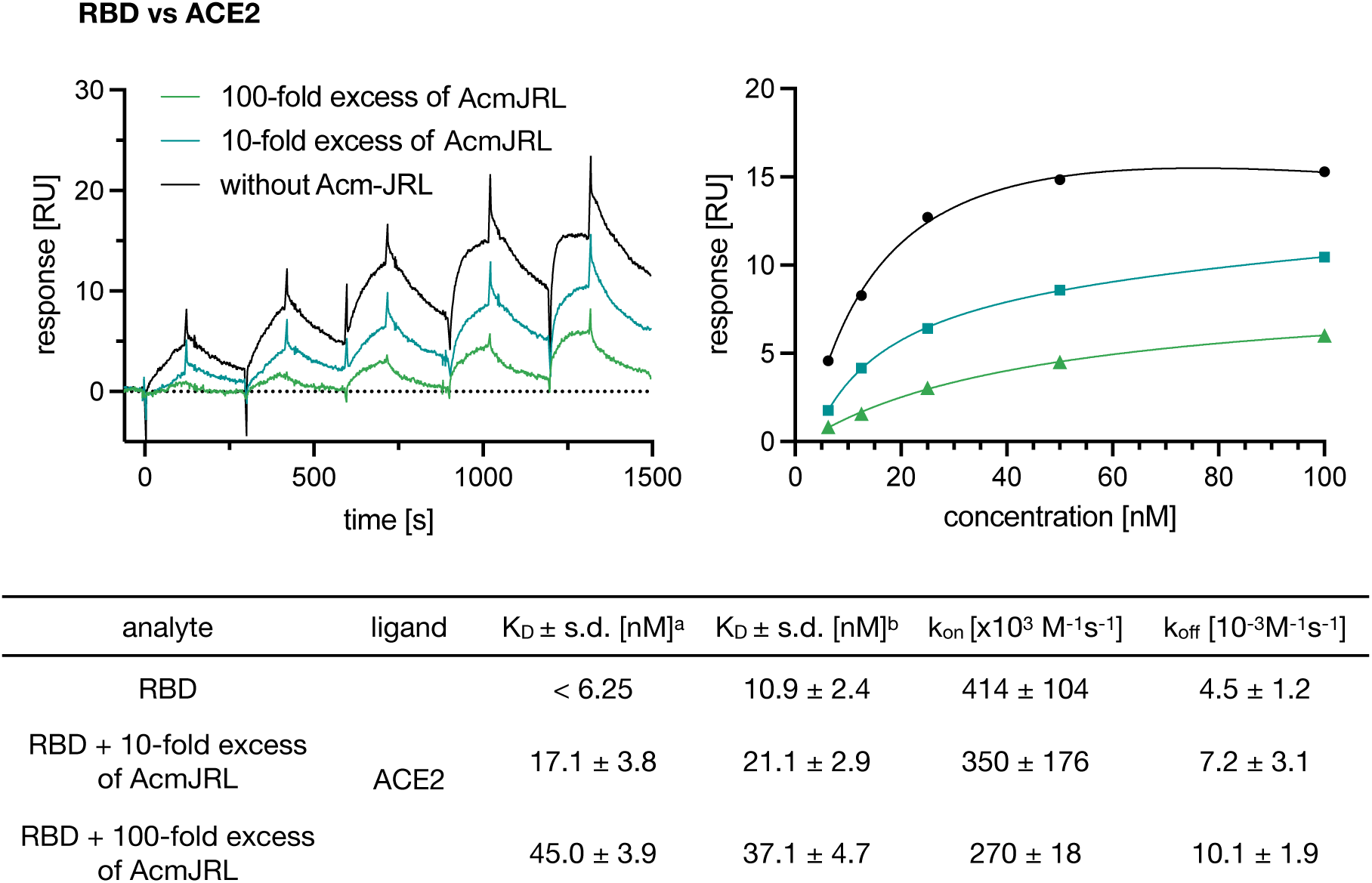
Surface plasmon resonance (SPR) sensorgrams and corresponding affinity plots for the binding of S-protein RBD against immobilised ACE2 (black). Prior to injection, S-RBD was also preincubated with 10-fold (blue) or 100-fold (green) excess of AcmJRL. Dissociation constants calculated from ^a^Langmuir isotherm or ^b^rate constants, K_D_= *k_off_*/*k_on_*.

The influence of AcmJRL on the affinity of S-protein RBD binding to the ACE2 receptor was studied by an addition of AcmJRL with a molar excess of lectin (factor 10 and 100, i.e. 1 μM and 10 μM Figure 8) to the RBD and preincubation prior to injection. Then, a single cycle kinetics run with injections of 5 dilutions (6.25 nM RBD with 10 and 100 fold excess AcmJRL – 100 nM RBD with 10 and 100 fold excess AcmJRL) of the RBD-AcmJRL mixture was performed with immobilised ACE2 receptor. Although residual binding of RBD to ACE2 receptor could still be observed (Figure 8), its apparent affinity was reduced by a factor of two (K_D_ = 21.1 ± 2.9 nM) after preincubation with a 10-fold excess of AcmJRL. An increase to a 100-fold excess of preincubated AcmJRL led to another twofold decrease in K_D_ value to 37.1 ± 4.7 nM. This rather moderate inhibitory effect probably resulted from the dilution of the preincubation mixture into the different injected concentrations, resulting in AcmJRL concentrations below the determined K_d_s of AcmJRL for spike RBD and ACE2. The addition of sufficiently high concentrations of AcmJRL to saturate RBD could not be used to overcome this problem, since AcmJRL also directly binds to the glycans of the immobilized ACE2 impacting on the recorded SPR response.

In a clinical scenario, saturating both the glycans of ACE2 and spike with AcmJRL could be beneficial to weaken the virus-host interaction and provide the immune system with an added advantage while battling the infection.

### Conclusion and Outlook

The current SARS-CoV-2 pandemic is a serious crisis that urgently asks for therapeutic treatment options. The viral envelope of SARS-CoV-2 is densely covered by the highly glycosylated spike protein, which is essential for viral cell entry via binding to the ACE2 receptor.

In this work, the pineapple-derived jacalin-related lectin AcmJRL was purified from the active pharmaceutical ingredient bromelain and characterised by mass spectrometry, differential scanning fluorimetry and dynamic light scattering. We further analysed the lectin’s ligand specificity by glycan array analysis using two complimentary arrays. The data further supported the previously reported preference of AcmJRL for mannopentaose. A solution phase binding assay was subsequently developed to quantify AcmJRL-carbohydrate interactions.

Then, the interaction of AcmJRL to recombinantly produced SARS-CoV-2 spike protein was studied by surface plasmon resonance analysis. The low μM binding was carbohydrate-dependent and could be inhibited by supplementation with mannotriose. Finally, we could show that addition of AcmJRL reduced the tight binding affinity of the spike RBD for the human ACE2 receptor. Thus, bromelain and specifically its component AcmJRL could constitute a novel antiviral drug to neutralise SARS-CoV-2 post exposure.

## Supporting information

CFG glycan array for AcmJRL

## Acknowledgements

The authors are thankful to Ursapharm (Saarbrücken, Germany) for funding parts of this work. We are indebted to Dr. Jamie Heimburg-Molinaro and Kelly Baker for the CFG glycan array analysis and we acknowledge the participation of the Protein-Glycan Interaction Resource of the CFG and the National Center for Functional Glycomics (NCFG) at Beth Israel Deaconess Medical Center, Harvard Medical School (supporting grant R24 GM137763). Furthermore, we would like to express our gratitude to Dr. Nadya Shilova from Semiotik (Moscow, Russia) for helpful suggestions for analyzing the Semiotik array. Further, we thank Alexandra Amann and Stefanie Neuber for performing cell culture experiments.

## Materials and Methods

Bromelain was supplied by Ursapharm (Saarbrücken, Germany). Mannotriose (> 95%), Me-α-L-Fuc (> 98%), Me-β-L-Fuc (> 98%) were obtained from Carbosynth Ltd (UK); D-glucosamine, D-mannose, D-galactose, D-Man-α-1,2-D-Man, D-Man-α-1,3-D-Man and D-Man-α-1,6-D-Man were obtained from Dextra Laboratories Ltd. (UK); Me-α-D-Glc (≥ 99%) and D-xylose (≥ 99) was obtained from Sigma-Aldrich Chemie GmbH (Germany); ; Me-β-D-Ara (> 99%) and maltose (> 98) were obtained from TCI Europe (Belgium); cellobiose (≥98%) was obtained from Carl Roth (Germany); L-rhamnose (> 99%) was obtained from AppliChem GmbH (Germany); melibiose was obtained from Alfa Aesar (UK); D-lyxose was obtained from Acros Organics (Belgium) and *N*-acetyl D-glucosamine was obtained from MP Biomedicals (France); raffinose (> 99%) was obtained from Th. Geyer GmbH & Co. KG (Germany); D-arabinose (> 98%) was obtained from Abcam (UK); L-fucose was obtained from Jennewein Biotechnologie GmbH (Germany).

### Preparation of D-mannosylated sepharose

D-Mannosylated sepharose was synthesised according to the protocol of Fornstedt and Porath(16): Sepharose CL-6B beads (Sigma-Aldrich Chemie GmbH, Germany, 15 mL) were suspended in Na_2_CO_3_-buffer (500 mM, pH 11, 15 mL). Divinylsulfone (1.5 mL) was added and the suspension was stirred at r.t. for 70 min. Activated sepharose was extensively washed with water and resuspended in 15 mL aqueous D-mannose solution (20% m/v, 500 mM Na_2_CO_3_, pH 10). The reaction was stirred over night at r.t., filtered and extensively washed with water. Unreacted activated sepharose was quenched by addition of β-mercaptoethanol (300 μL) in buffer (15 mL, 500 mM NaHCO_3_, pH 8.5) for 120 min. After filtration and washing of the mannosylated beads, they were filled into 5 mL plastic columns for affinity chromatography.

### Isolation of AcmJRL from bromelain

AcmJRL was isolated in analogy to the protocol of Azarkan *et al*.^(12)^. Bromelain powder (28 g) was suspended in an Erlenmeyer flask in buffer (400 mL, 100 mM NaOAc pH 5, 1 mM EDTA, 20 mM methyl methanethiosulfonate) and stirred for 60 min at r.t.. After centrifugation (30,000 rcf, 30 min, 4 °C), the supernatant was dialysed twice for 1 h against 4 L Tris-buffered saline (TBS: 150 mM NaCl, 50 mM Tris pH 7.4). The sample was loaded on a D-mannosyl-sepharose column pre-equilibrated with the dialysis buffer. After extensive washing, the lectin was eluted with 1 M D-mannose in buffer. Eluted fractions containing AcmJRL were pooled and dialysed against TBS (5 × 3 h against 2 L). The yield (31 mg) was determined by UV-absorption at 280 nm (MW = 15.34 kDa, ε = 19940 M^−1^ × cm^−1^).

### Intact protein mass determination

Intact protein mass measurements for AcmJRL were performed on a Dionex Ultimate 3000 RSLC system using an Aeris Widepore XB C8, 150 × 2.1 mm, 3.6 μm dp column (Phenomenex, USA). Separation of a 2 μL sample was achieved by a linear gradient from (A) H_2_O + 0.1% formic acid to (B) MeCN + 0.1% formic acid at a flow rate of 300 μL/min and 45 °C. The gradient was initiated by a 1 min isocratic step at 2% B, followed by a linear increase to 75% B in 10 min to end up with a 3 min step at 75% B before re-equilibration with initial conditions. UV spectra were recorded on a DAD in the range from 200 to 600 nm. The LC flow was split to 37.5 μL/min before entering the maXis 4G hr - ToF mass spectrometer (Bruker Daltonics, Bremen, Germany) using the standard Bruker ESI source. In the source region, the temperature was set to 200 °C, the capillary voltage was 4000 V, the dry-gas flow was 5.0 L/min and the nebuliser was set to 1.0 bar. Mass spectra were acquired in positive ionisation mode ranging from 150 – 2500 *m/z* at 2.0 Hz scan rate. Protein masses were deconvoluted by using the Maximum Entropy algorithm (Spectrum Square Associates, Inc.).

### Dynamic light scattering

Dynamic light scattering (DLS) measurements were performed on a Zetasizer Nano-ZS (MalvernInstruments, UK). Protein solutions were filtered with a syringe filter (0.45 μm) before measurements. AcmJRL (14 μM) was measured in TBS (150 mM NaCl, 50 mM Tris pH 7.4) at 25 °C.

### Differential scanning fluorimetry

20 μL of a solution containing AcmJRL (20 μM) and SyproOrange (final concentration 10x of a 5000x stock in DMSO, Sigma-Aldrich, Germany) in TBS (150 mM NaCl, 50 mM Tris pH 7.4) was added to a white semi-skirted 96-well plate (Thermo Fisher) in triplicates. The melting curve measurements (25 – 95 °C, 0.5 °C/min) were performed and analysed on a real time PCR instrument (StepOnePlus, Applied Biosystems).

### Direct binding of fluorescent mannoside ligand 1 to AcmJRL

Fluorescent ligand **1**(24) was dissolved in DMSO, then diluted to a final concentration of 200 nM in TBS (150 mM NaCl, 50 mM Tris pH 7.4). AcmJRL was concentrated (Vivaspin column, 10,000 MWCO, Sartorius Stedim Biotech GmbH, Germany) to a concentration of 1.33 mM in TBS. After centrifugation (10 min, 25,000 rcf, r.t.), the concentration of AcmJRL was adjusted to 1 mM as determined by UV absorbance measurement at 280 nm (ε = 19,940 M^−1^ cm^−1^). A serial dilution of AcmJRL was dispensed in triplicates (10 μL each) in a black 384-well plate (Greiner Bio-One, Germany, cat no 781900). The solution of the fluorescent ligand (10 μL) was added to yield a final concentration of 100 nM. After incubation for 1 h at r.t., fluorescence (excitation 485 nm, emission 535 nm) was measured in parallel and perpendicular to the excitation plane on a PheraStar FS plate reader (BMG Labtech GmbH, Germany) and polarization was calculated. The data were analysed using a four-parameter fit calculated with MARS Data Analysis Software (BMG Labtech GmbH, Germany). Three independent measurements on three plates was performed.

### Reporter ligand displacement assay

The assay was performed in analogy to the protocol from Joachim *et al*. (21): A serial dilution of the test compounds was prepared in TBS (150 mM NaCl, 50 mM Tris pH 7.4). For single point inhibition measurements, carbohydrates were dissolved in TBS at 20 mM. A concentrated solution of AcmJRL was diluted in TBS together with the fluorescent reporter ligand **1** to yield concentrations of 40 μM protein and 20 or 200 nM ligand, respectively. 10 μL of this mix was added to 10 μL previously prepared dilutions of the test compounds in black 384-well microtiter plates (Greiner Bio-One, Germany, cat. no. 781900) in triplicate. After centrifugation (2680 rcf, 1 min, r.t.), the reactions were incubated for 60 min at r.t. in a humidity chamber. Fluorescence (excitation 485 nm, emission 535 nm) was measured in parallel and perpendicular to the excitation plane on a PheraStar FS plate reader (BMG Labtech GmbH, Germany). The measured intensities were reduced by the values of only AcmJRL in TBS, and fluorescence polarisation was calculated. The data were analysed with the MARS Data Analysis Software and fitted according to the four-parameter variable slope model. Bottom and top plateaus were fixed according to the value in absence of inhibitor and to the highest concentration of mannoside, respectively, and the data was reanalysed with these values fixed.

### Fluorescence labelling of AcmJRL

*FITC*: AcmJRL was diluted in Na_2_CO_3_-buffer (100 mM, pH 9.3) and concentrated (Vivaspin, Sartorius Stedim Biotech GmbH, 10,000 MWCO) to yield a final protein concentration of 78 μM (1.8 mg in 1.5 mL). FITC (Merck, Germany, 95 μL of a freshly prepared 7.7 mM solution in carbonate buffer pH 9.3, 0.73 μmol, 6.2 eq.) was added and incubated for 1 h at r.t.. The reaction was quenched with ethanolamine (1 μmol, 8.3 eq.) for 1 h at r.t.. Excess reagents were removed by filtration (Vivaspin, 10000 MWCO), then the protein was affinity-purified as described above for unlabelled AcmJRL. The protein concentration and degree of labelling (DOL) was calculated according to the manufacturers protocol (Thermo Scientific, Rockford, USA):

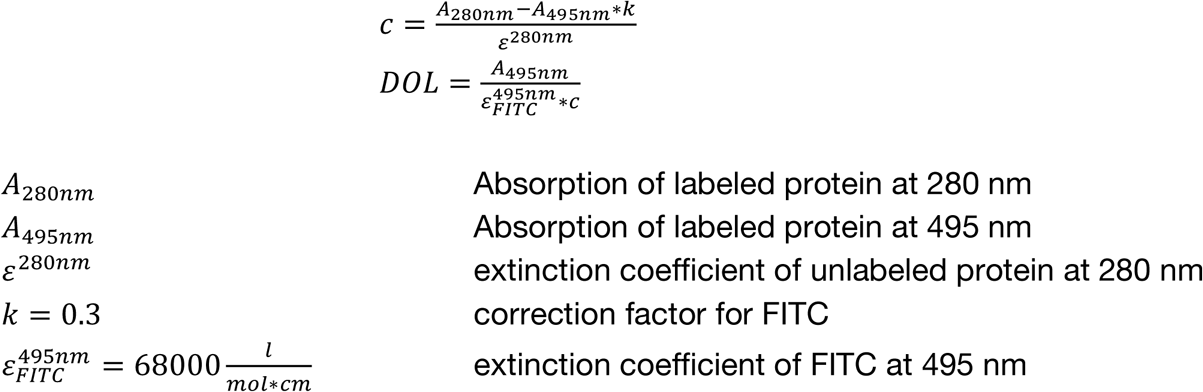

#### NHS-activated Cy3

AcmJRL was diluted in PBS pH 8.4 and concentrated (Vivaspin, Sartorius Stedim Biotech GmbH, 10,000 MWCO) to yield a final protein concentration of 293 μM (4.5 mg in 1.5 mL). NHS-activated Cy3 (Lumiprobe, Germany, 75 μL of a freshly prepared 29 mM solution in DMSO, 2.2 μmol, 9 eq.) was added and incubated for 5 h at r.t.. Excess reagents were removed by filtration (Vivaspin, Sartorius Stedim Biotech GmbH, 10000 MWCO), then the protein was affinity-purified as described above for unlabelled AcmJRL. The protein concentration and degree of labelling (DOL) was calculated as described above.

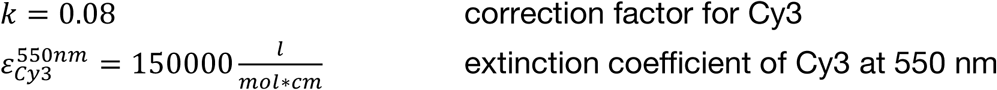

### Glycan array analysis

FITC-labeled AcmJRL was tested by the National Center for Functional Glycomics (NCFG, Boston, MA, USA) on the CFG glycan microarray version 5.5 containing 585 printed glycans in replicates of 6. Standard procedures of NCFG (details see https://ncfg.hms.harvard.edu/files/ncfg/files/protocol-direct_glycan_binding_assay-cfg_slides.docx) were run with 5 and 50 μg/mL protein based on the protocol by Blixt et al. (19). Raw-data (Tables S5 and S6) will be shared online on the CFG website. Cy3-labelled AcmJRL was tested in-house on a glycan microarray slide from Semiotik LLC (Moscow, Russia) containing 610 printed glycans in replicates of 6. Standard procedures were run at 20, 200 and 400 μg/mL based on the protocol by Olivera-Ardid *et al*.(20). Fluorescence intensity was measured at 565 nm upon excitation at 520 nm on a Sapphire Biomolecular imager (Azure Biosystems, Dublin, CA, USA) at 10 μm resolution. Scan data was processed with ScanArray software (Perkin Elmer, Waltham, MA, USA), using OSPS090418_full.360.80 um.gal (kindly provided by Semiotik) for dot-glycan assignment. Raw data (dot mean fluorescence intensity) was processed by GraphPad Prism 9 (GraphPad Software, USA). Processed data in table format is provided in the appendix.

### Protease activity assay

Purified AcmJRL was used and adjusted to A_280nm_ = 1. For the positive control, 2 mg bromelain powder were suspended in 2 mL buffer (100 mM KCl, 5 mM KOAc, 5 mM HOAc, pH 4.6) and incubated at 37 °C, 500 rpm on a Thermomixer (Eppendorf, Germany) for 10 min. After centrifugation (17600 rcf, 10 min, r.t.) the supernatant adjusted to A_280nm_ = 1. Then, 4 μL protein or bromelain extract was diluted in 200 μL buffer. Z-Lys-ONP (Merck, Germany, 4 μL from a 25 mM solution in H_2_O/MeCN 1:1) was added and absorption at 340 nm was measured on a CLARIOstar microplate reader (BMG Labtech, Germany) for 300 s (5 s interval). Blank controls without added protein/bromelain and substrate were performed for background subtraction. Data was analysed by using GraphPad Prism 9 (GraphPad Software, USA).

### Cloning and Recombinant Expression of SARS-CoV-2 spike protein

A synthetic DNA fragment was purchased from Eurofins MWG. The nucleotide sequence was codon optimised for mammalian cell expression (translational amino acid sequence based on PDB code: 6VXX (29)). The nucleotide sequence coding for the extraviral domain of SARS-CoV-2 spike (coding for aa 1-1213) was amplified via PCR with the restriction sites 5′-BamHI/XhoI-3′ (Fw Primer: ATAT<u>GGATCC</u>ATGTTCGTGTTCCTGGTTCTT; Rv Primer: AATATGAGCAGTACATAAAATGGCCC<u>CTCGAG</u>ATAT; purchased from Merck). As vector system, the in house vector πα-SHP-H (provided by Dr. Jesko Köhnke, pCAGGS based, NCBI accession number: LT727518) was chosen for the mammalian expression system. The amplicon was digested with the respective restriction enzymes (ThermoFisher) and ligated with T4 Ligase (ThermoFisher) into the linearised πα-SHP-H vector to clone πα-SHP-H–Sgene with an N-terminal octahistidin tag.

The mammalian cell line HEK 293/T served as host for recombinant production of the glycoprotein. The cells were cultured with Dulbecco’s Modified Eagle Medium (DMEM, Sigma Aldrich) in a Hyperflask M (A= 1720 cm^2^, Corning) and incubated at 37 °C under 5% CO_2_ atmosphere until a confluence of 80-90% was reached. The transient mammalian transfection was applied in a mass ratio of 1:2 πα-SHP-H-Sgene : PEI (linear, MW > 25000, Alfa Aesar) (30). After 5 h incubation, the medium was exchanged and the transfected cell line was incubated for 48 h at 37 °C under 5% CO_2_ atmosphere before the supernatant was filtered (0.22 μm pore size) and stored on ice. The supernatant was loaded onto a 5 mL HisTrap HP column and washed with 20 column volumes lysis buffer (200 mM NaCl, 20 mM Tris, 20 mM imidazol pH 8) and the protein was obtained with elution buffer (200 mM NaCl, 20 mM Tris, 500 mM imidazole pH 8). Size exclusion chromatography was performed on a HiLoad 16/600 Superdex 200 PG. The elution volume of 60-70 mL was collected and the solution was concentrated to a volume of 200 μL with an Amicon Ultra-15 PLHK Ultracel-PL Membrane 100 kDa (Merck) spin column. The amount of SARS-CoV-2 spike protein was determined via UV absorbance measurement on a Nanodrop 2000c to a mass concentration of 7.1 mg/mL (ε = 141,35* 10^3^ L·mol^−1^·cm^−1^; MW = 142.13 kDa). For SDS-PAGE, 20 μL (6 μg) of SARS-CoV-2 spike protein (in 150 mM NaCl, 20mM HEPES pH 8) was mixed with 4 μL 6x SDS-loading buffer (1.2 g SDS, 6 mg bromophenolblue, 4 mL glycerol, 0.6 mL 1 M Tris pH 8.0, 5.4 mL H_2_O, 930 mg dithiothreitol) and heated for 2 min at 100 °C, afterwards cooled to r.t.. A SDS-PAGE gel (10% v/v polyacrylamide) was loaded with 20 μL of the prepared sample. 6 μL PageRulerPlus Prestained Protein Ladder (ThermoFisher Scientific) was loaded to monitor the progress of the SDS-PAGE and to estimate the approximate size of the protein after staining the gel. The SDS-PAGE was run in a Mini-PROTEAN Tetra System (BIO RAD) with SDS Laemmli buffer at 150 V for 100 min. The gel was stained with Coomassie Brilliant Blue.

### Deglycosylation with PNGase F

To 5 μg of spike-protein in 20 μL of 50 mM sodium phosphate buffer (pH 7.5), 2 μL (1000 U) PNGase F (New England Biolabs Inc) were added and the reaction was incubated for 6 h at 37 °C. As a negative control, the same amount of spike-protein was incubated in absence of enzyme in otherwise identical conditions.

### Surface Plasmon Resonance (SPR)

SPR experiments were conducted on a Biacore X100-system (GE Healthcare). SARS-CoV-2 spike-protein was immobilised on a CM5 sensorchip after EDC/NHS activation via amine coupling (0.5 M 1-ethyl-3-(3-dimethylamino-propyl) carbodiimide hydrochloride and 0.1 M N-hydroxysuccinimide in water for activation). The glycoprotein was dissolved in 10 mM sodium acetate pH 4.5 (50 μg/mL) and injected over the activated chip surface (contact time of 200 s, flow rate of 30 μL/min), followed by ethanolamine for deactivation of excess reactive NHS ester groups in to obtain a final response of ∼6700 RU. The ACE2 receptor (Sigma Aldrich, SAE0064) was immobilised on a CM5 sensorchip using the same conditions and a final response of ∼990 RU was obtained. The SARS-CoV-2 spike RBD (ThermoFisher) was immobilised on a CM5 sensorchip under the same conditions and a final response of ∼1970 RU was obtained. SPR assays were performed with a flow rate of 30 μL min in HBS-EP (150 mM NaCl, 10 mM HEPES pH 7.4, 3 mM EDTA, 0.005% Polysorbate-20). Increasing concentrations of AcmJRL (1.25, 2.5, 5.0, 10.0, 20 μM) with or without 10 mM mannotriose in the sample buffer were injected for single cycle measurements (120 s contact time, 180 s dissociation time). Respectively, increasing concentrations of RBD (6.75 – 100 nM) with or without AcmJRL in a molar ratio of RBD:AcmJRL of 1:10 and 1:100 were injected for single cycle measurements (120 s contact time, 180 s dissociation time). The CM5 chip surface was regenerated by one injection of ethylene glycol 80% v/v (contact time 30 sec, flow rate 30 μL/ min, followed by 2 × injections of HBS-EP (60 sec, flow rate 30μL/min). Binding analysis results were evaluated using the Biacore X100 evaluation software to plot the sensorgrams and the Langmuir graphs, and to determine dissociation constants. Graphs were visualised with Graphpad Prism 9 software (Graph Pad Software, San Diego, CA, USA). Spikes caused by instrumental effects were removed by omitting 3-5 seconds around each injection.

### Supporting Information

Additional data can be found in the supporting information pdf file. Raw data for the CFG Glycan Array for AcmJRL can be found in the attached xls file.

