## Supporting Information for "The Glycan-Specificity of the Pineapple Lectin AcmJRL and its Carbohydrate-Dependent Binding of the SARS-CoV-2 Spike Protein"

**Table of contents**

| Protein glycation (Figure S1) | S1 |
| --- | --- |
| DSF of AcmJRL and three-dimensional dimensional structure (Figure S2) | S1 |
| Residual protease activity screen (Figure S3) | S2 - 3 |
| Accessible lysines on the protein surface of AcmJRL (Figure S4) | S3 |
| Row number assignment for Semiotic glycan ID (Table S1) | S4 - S12 |
| Raw data from Semiotic glycan array (Table S2) | S13 -S17 |
| Trypsin digest of SARS-CoV-2 spike-protein | S18 |
| Sample preparation for MALDI-MS | S18 |
| MALDI-MS | S18 |
| Spectra and peak list processing | S18 |
| Mass spectrum of SARS-CoV-2 spike after trypsin digest (Figure S5) | S19 |
| Parameter input into post processing software (Table S3) | S19 |
| Mass list with corresponding intensity (Table S4) | S20 |
| Primary sequence of SARS-CoV-2 spike protein | S21 |

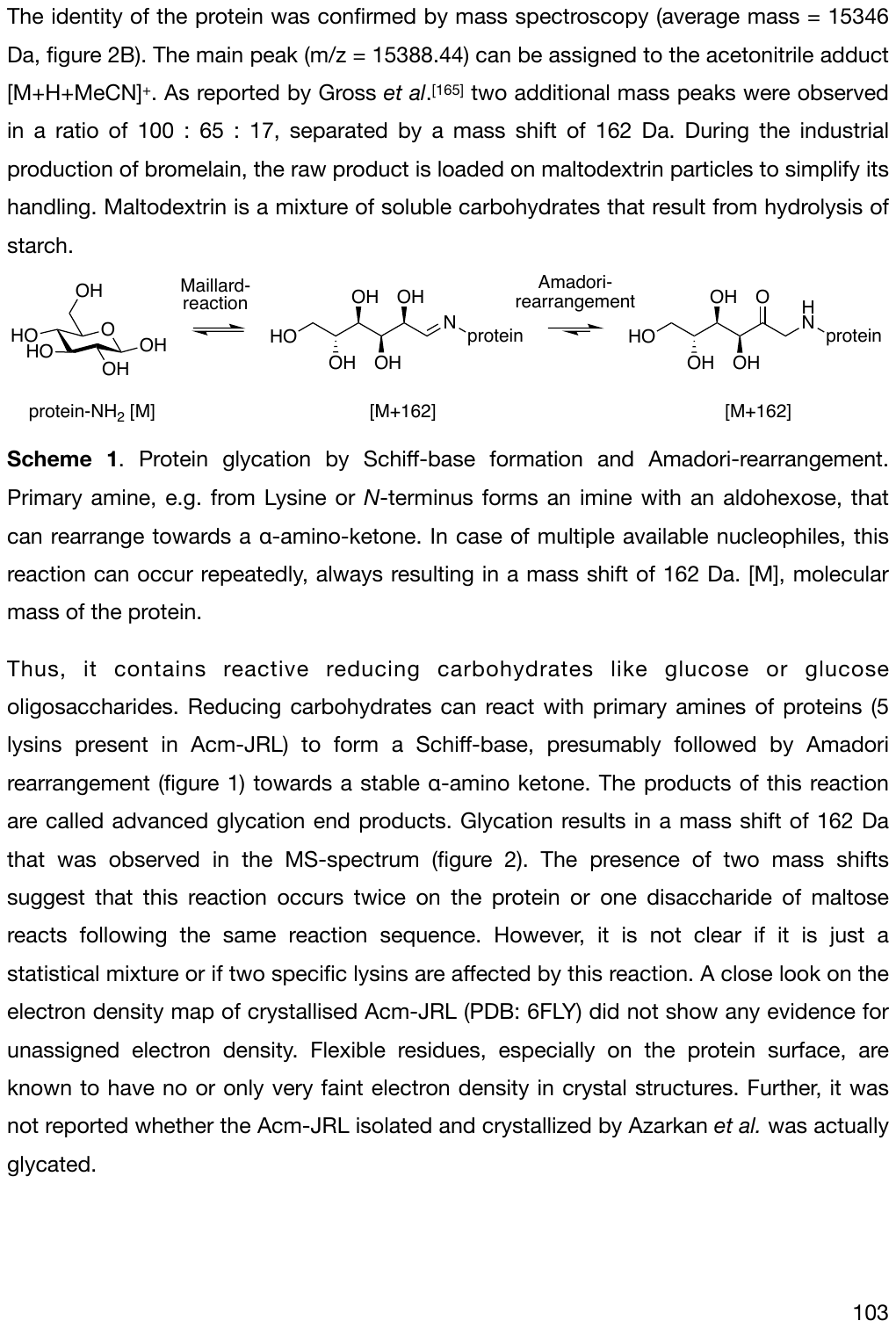

**Figure S1**. Protein glycation by Schiff-base formation and Amadori-rearrangement. Primary amine, e.g. from lysine or *N*-terminus forms an imine with an aldohexose, that can rearrange towards a α-amino-ketone. In case of multiple available nucleophiles, this reaction can occur repeatedly, always resulting in a mass shift of 162 Da. [M], molecular mass of the protein.

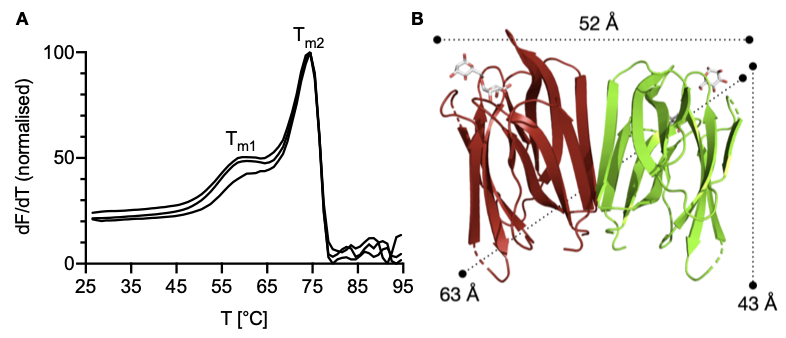
**Figure S2**. **A** Differential scanning fluorimetry of AcmJRL: biphasic denaturation together with two peaks Tm1 and Tm2 correspond to the reported dimeric structure in solution. **B** Global structure and dimensions of AcmJRL in the crystal structure (PDB: 6FLY).

**Residual protease activity screen**

Protease impurities from the soluble protein fraction of bromelain can disturb future experiments and thus have to be quantitatively removed. No significant proteolytic activity was detected following purification of AcmJRL (figure S3).

N^α^-benzyloxycarbonyl-l-lysine o-nitrophenyl ester (Z-Lys-ONp) was used as a model substrate that releases chromogenic o-nitrophenol (HONp) upon proteolysis.

After linear regression fit of the raw data (R^2^ > 0.90), the slope was used to describe the proteolytic activity (figure S3). Unpurified soluble protein fraction of bromelain led to a fast release of HONp (slope^-1^ = 542 s). In contrast, purified lectin showed only very little release of HONp (slope^-1^ = 8984 s), comparable to the absence of protein (negative control, slope^-1^ = 9430 s). In conclusion, AcmJRL was obtained in good purity and can be used for further studies.

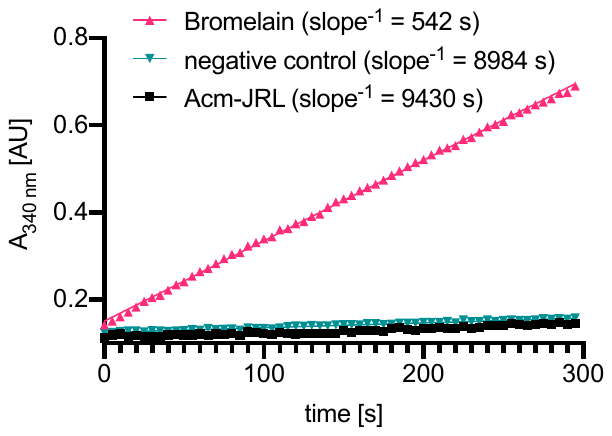

**Figure S3**. Protease activity of bromelain before and of AcmJRL after purification by affinity chromatography. Negligible proteolytic activity was observed for the purified protein. Soluble protein fraction from bromelain and absence of protein were used as positive and negative control, respectively.

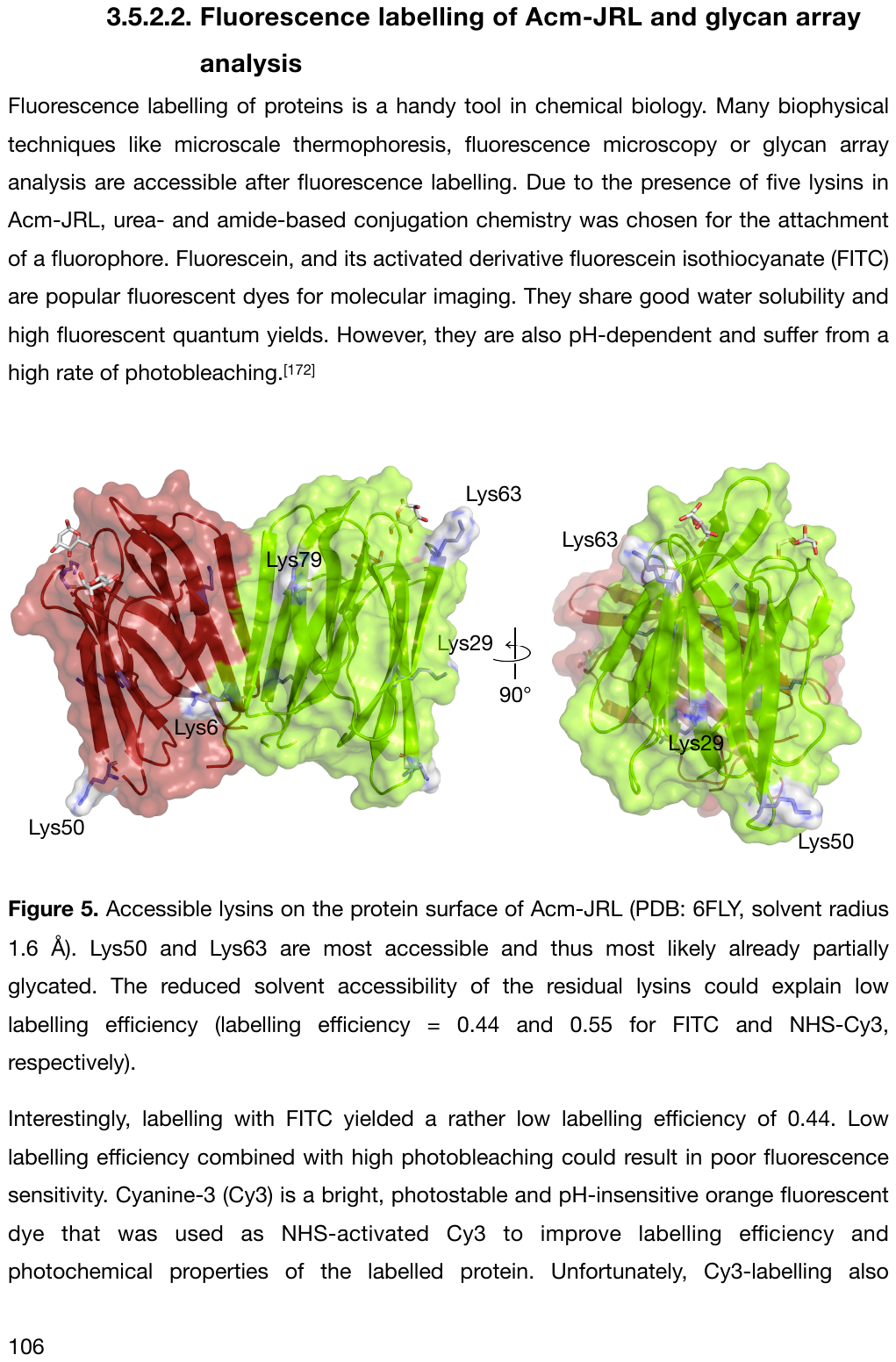

**Figure S4.** Accessible lysines on the protein surface of AcmJRL (PDB: 6FLY, solvent radius 1.6 Å). Lys50 and Lys63 are most accessible and thus most likely already partially glycated. The reduced solvent accessibility of the residual lysines could explain low labelling efficiency (labelling efficiency = 0.44 and 0.55 for FITC and NHS-Cy3, respectively).

**Table S1**. Row number assignment for Semiotic glycan ID.

saccharide nomenclature, row number, SGID

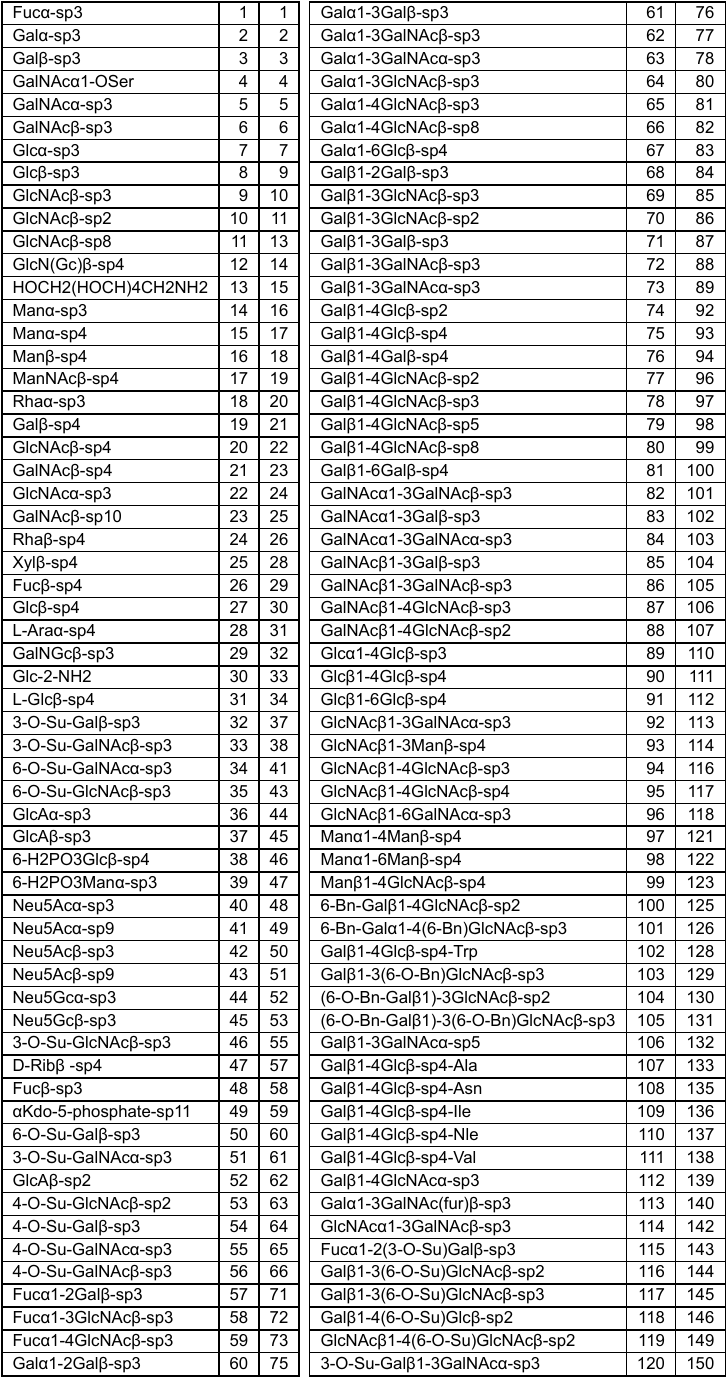

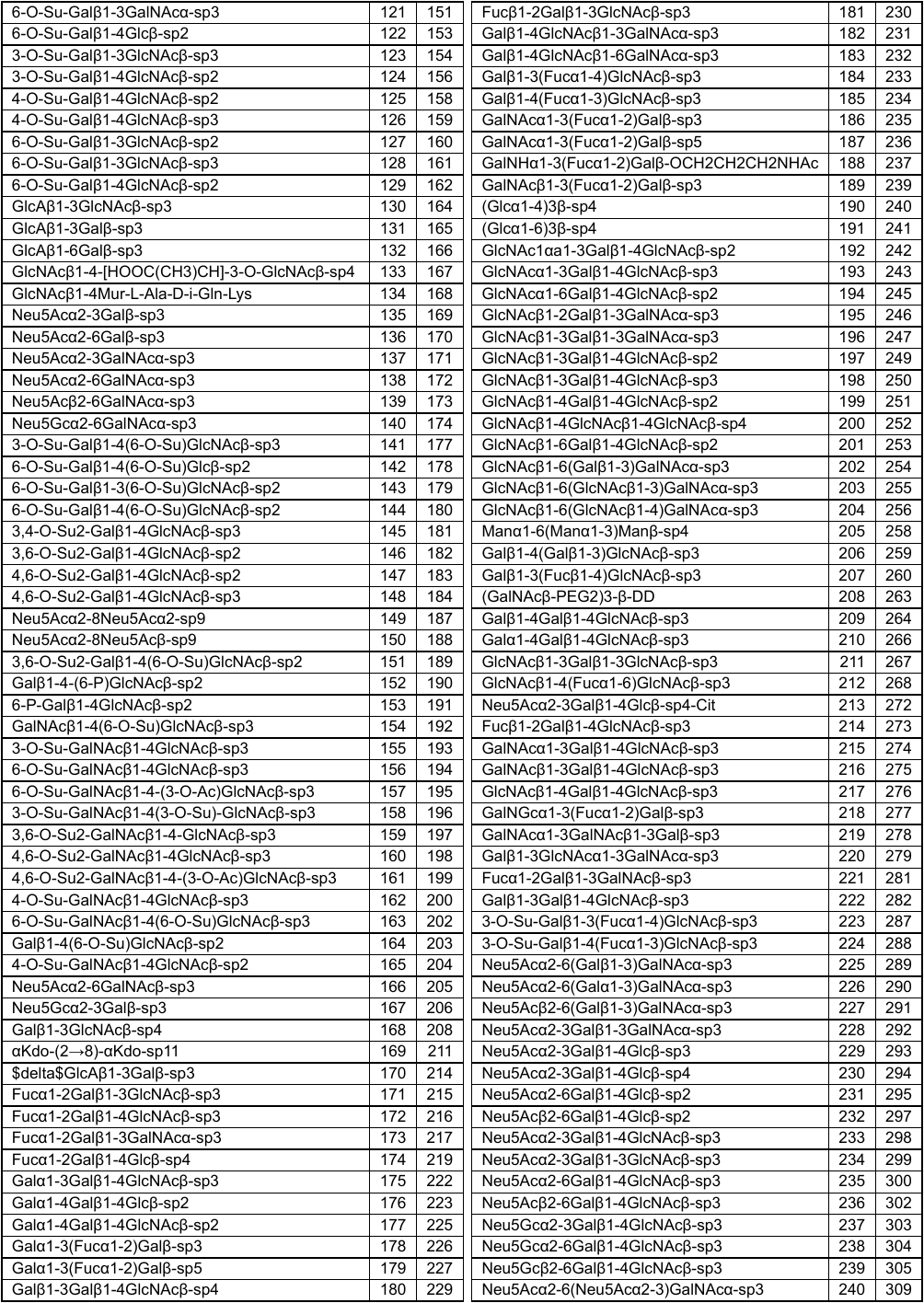

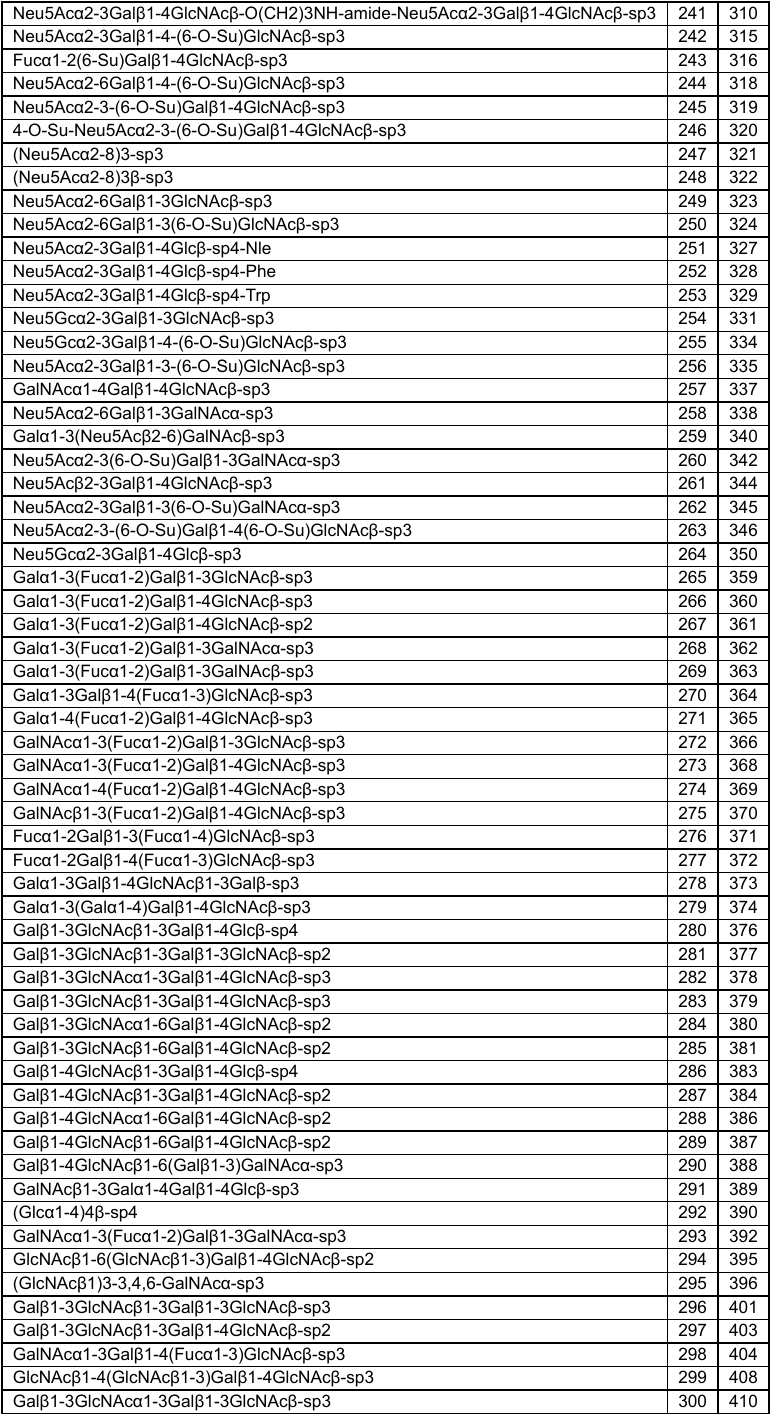

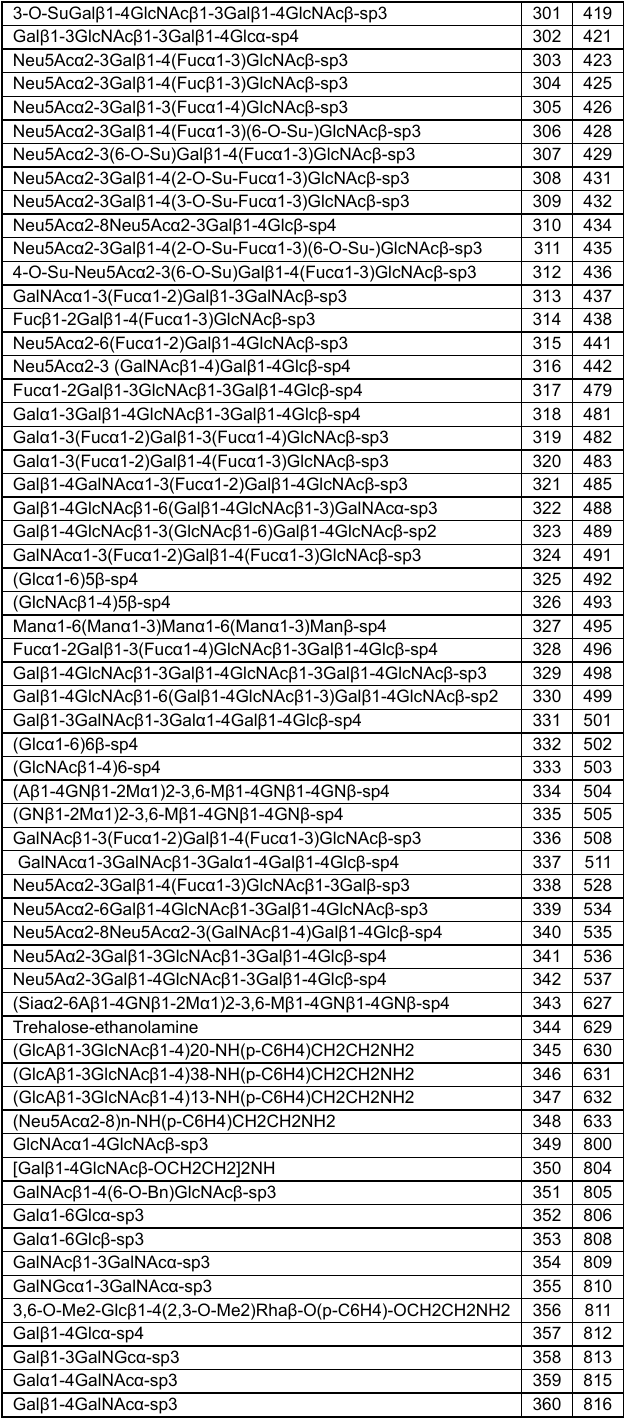

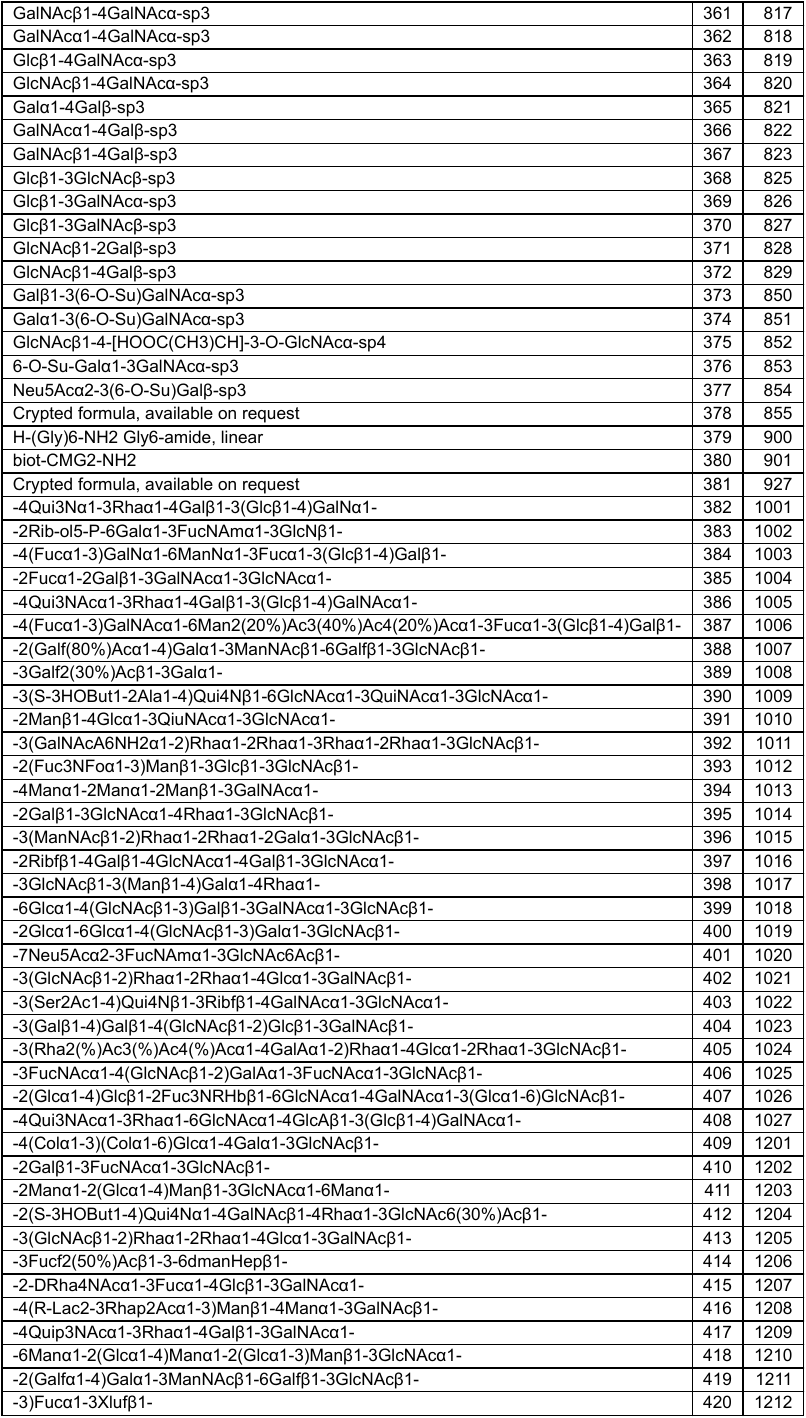

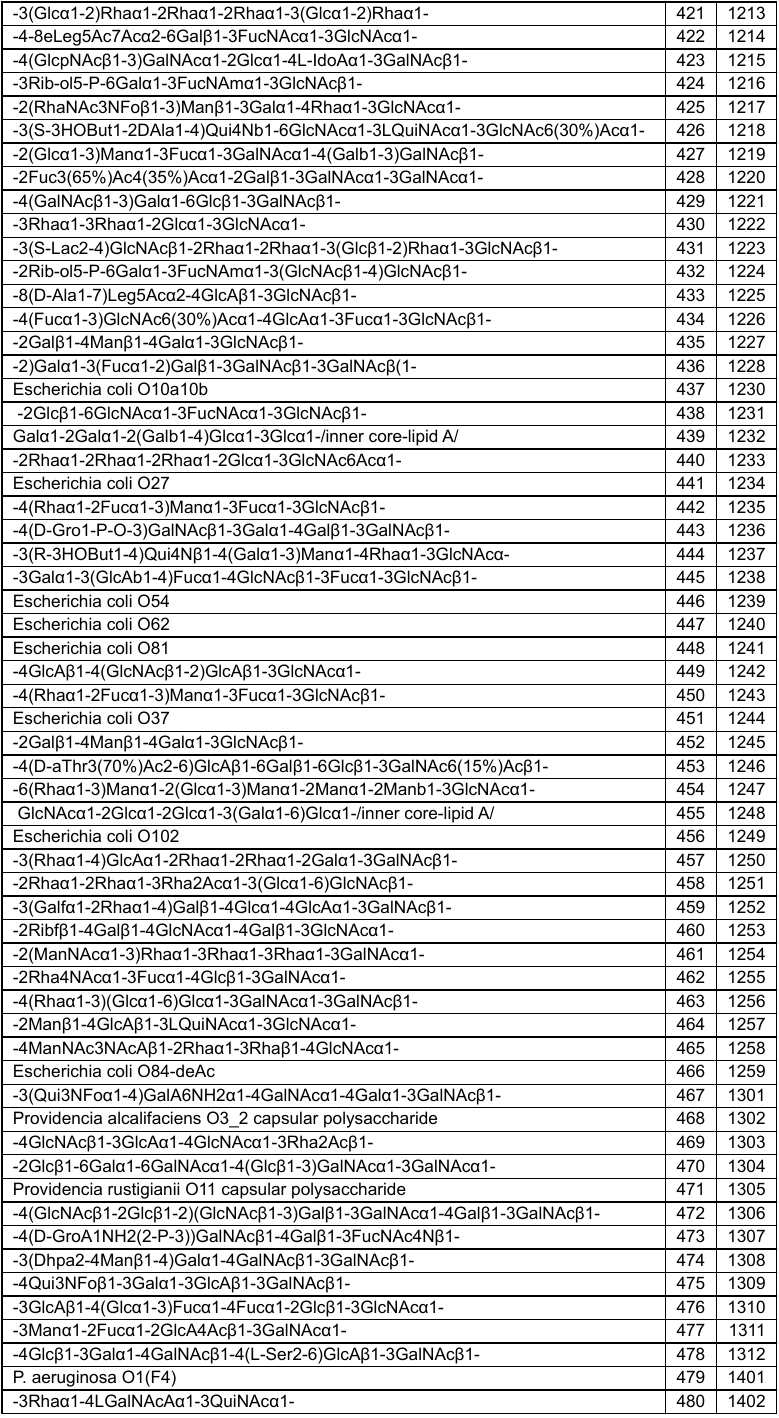

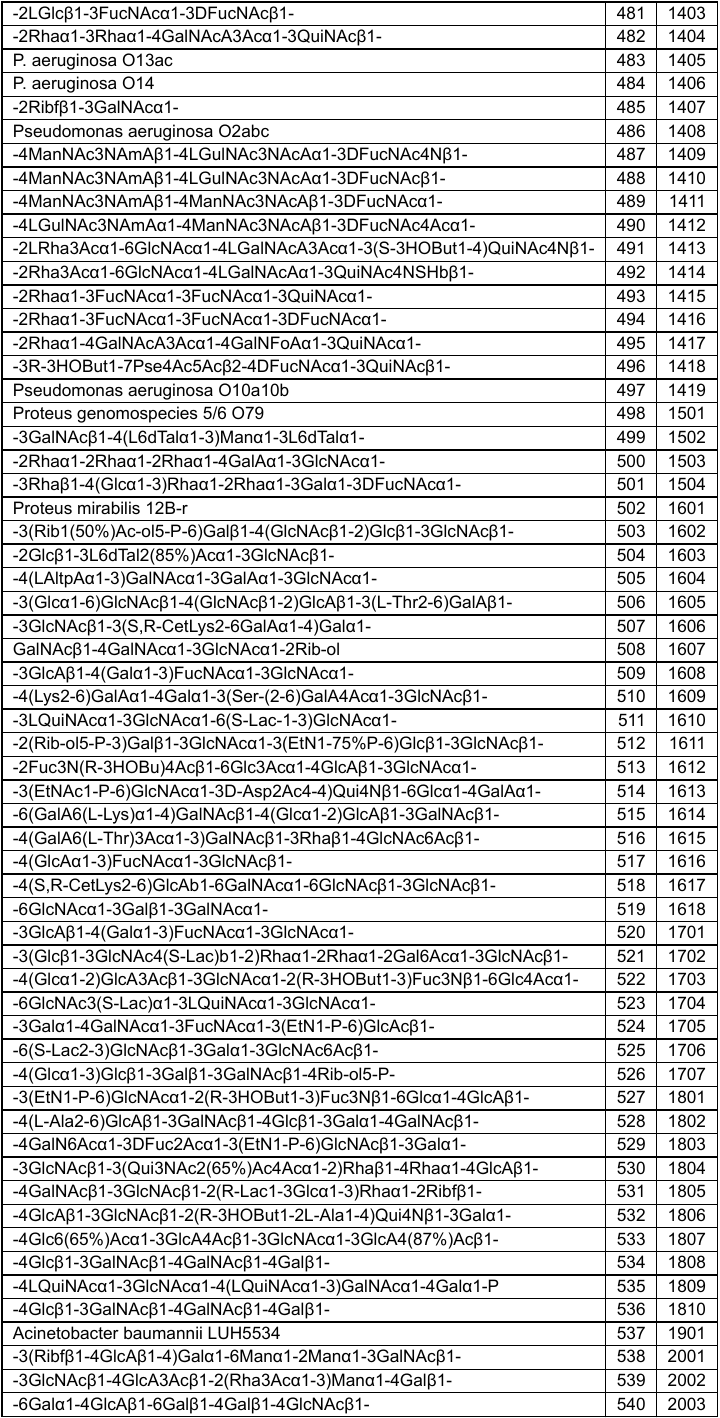

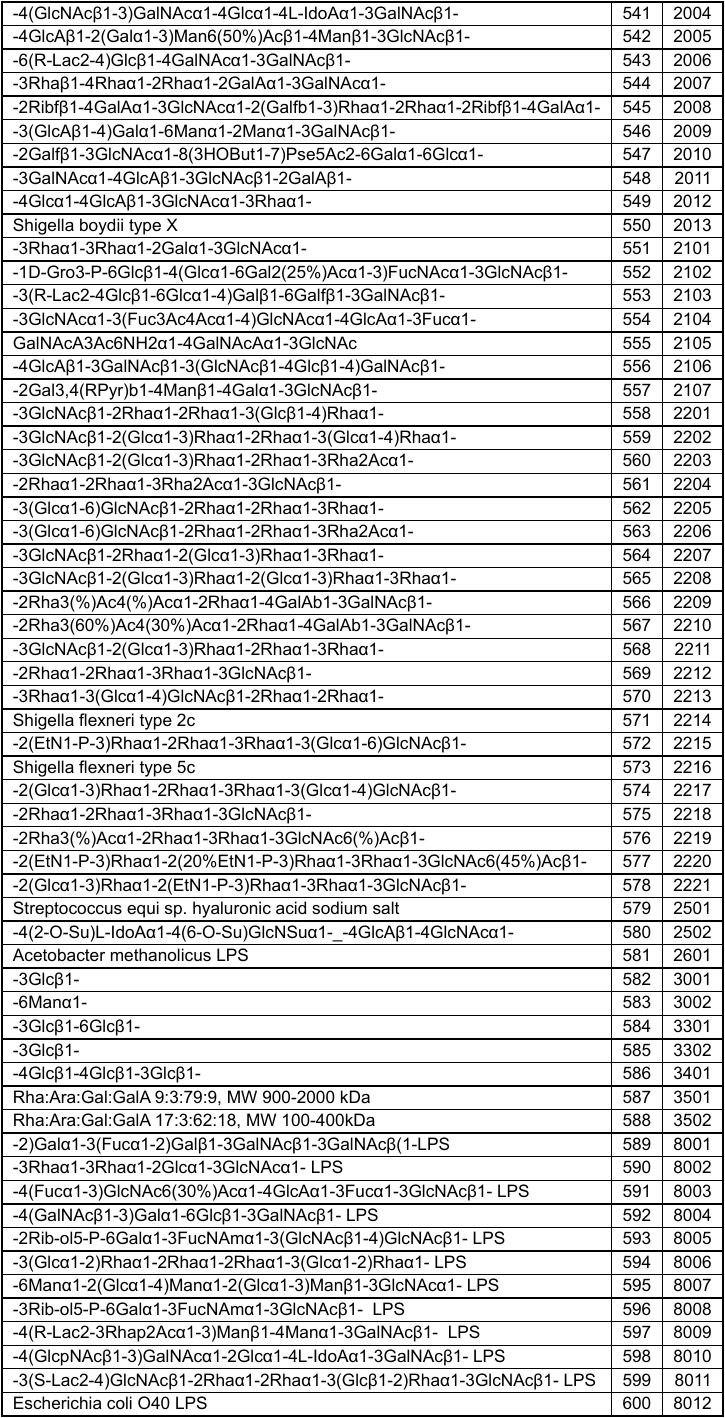

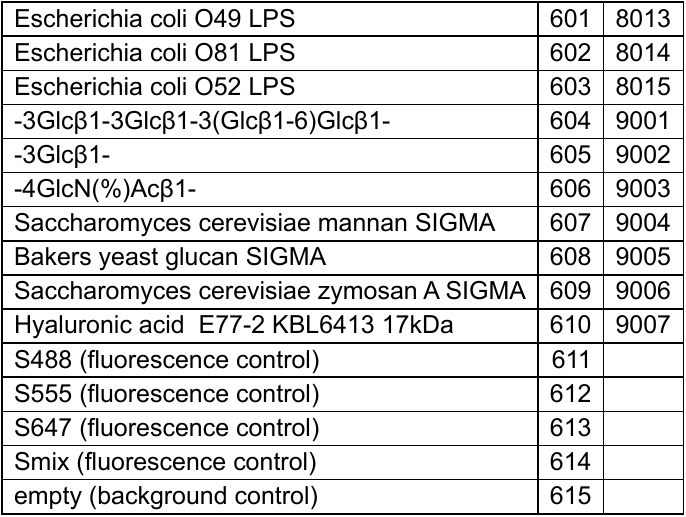

**Table S2**. Raw data from Semiotic glycan array.

columns correspond to: trivial name, row number, Semiotik glycan ID, mean fluorescence intensity, s.d.

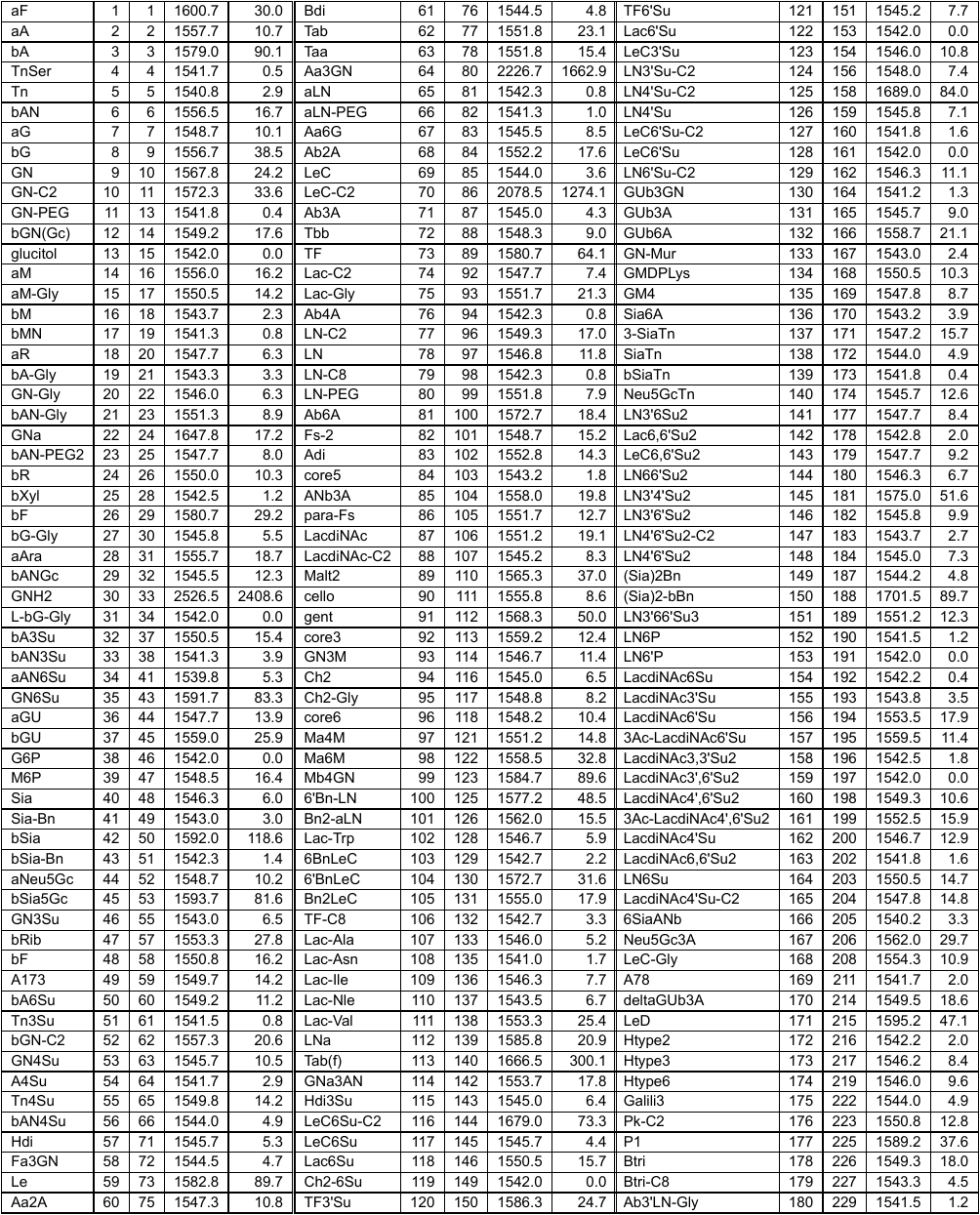

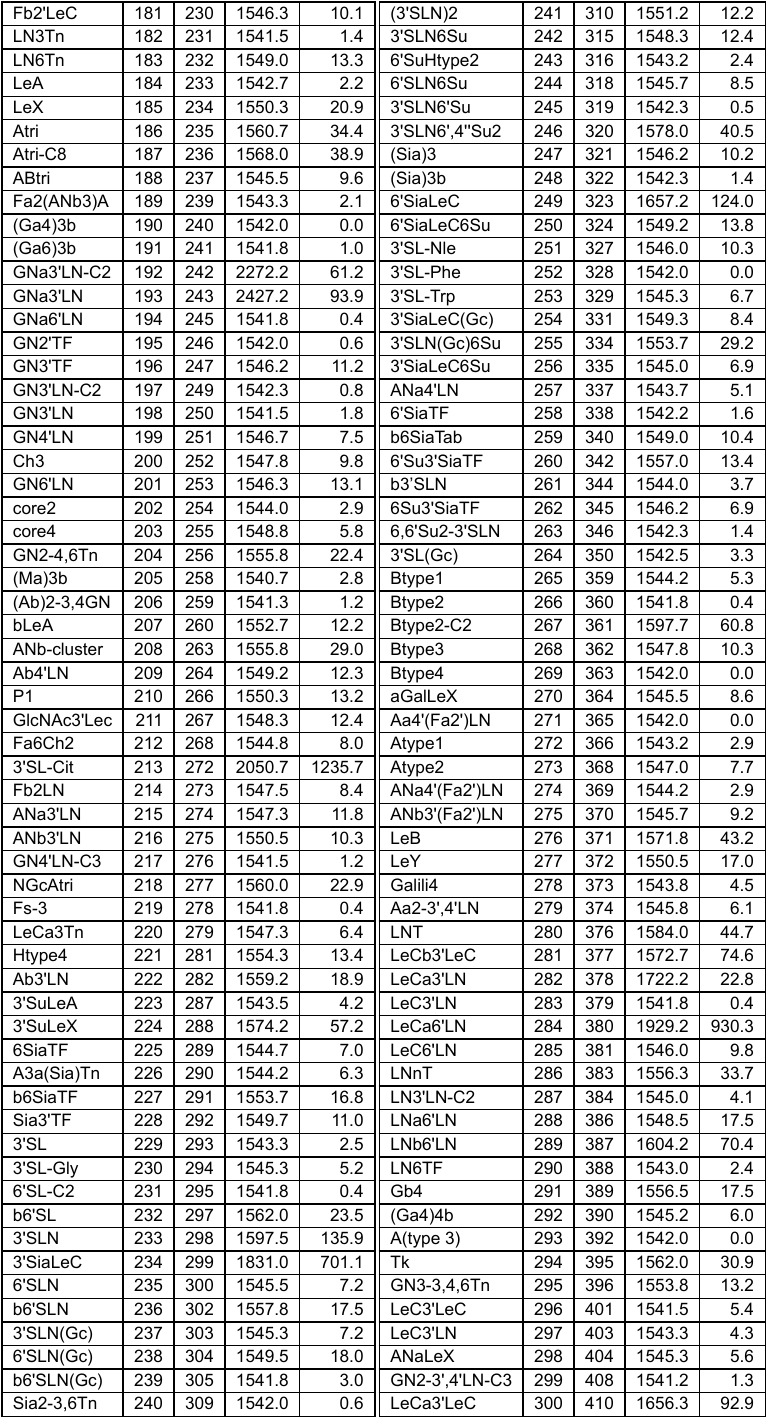

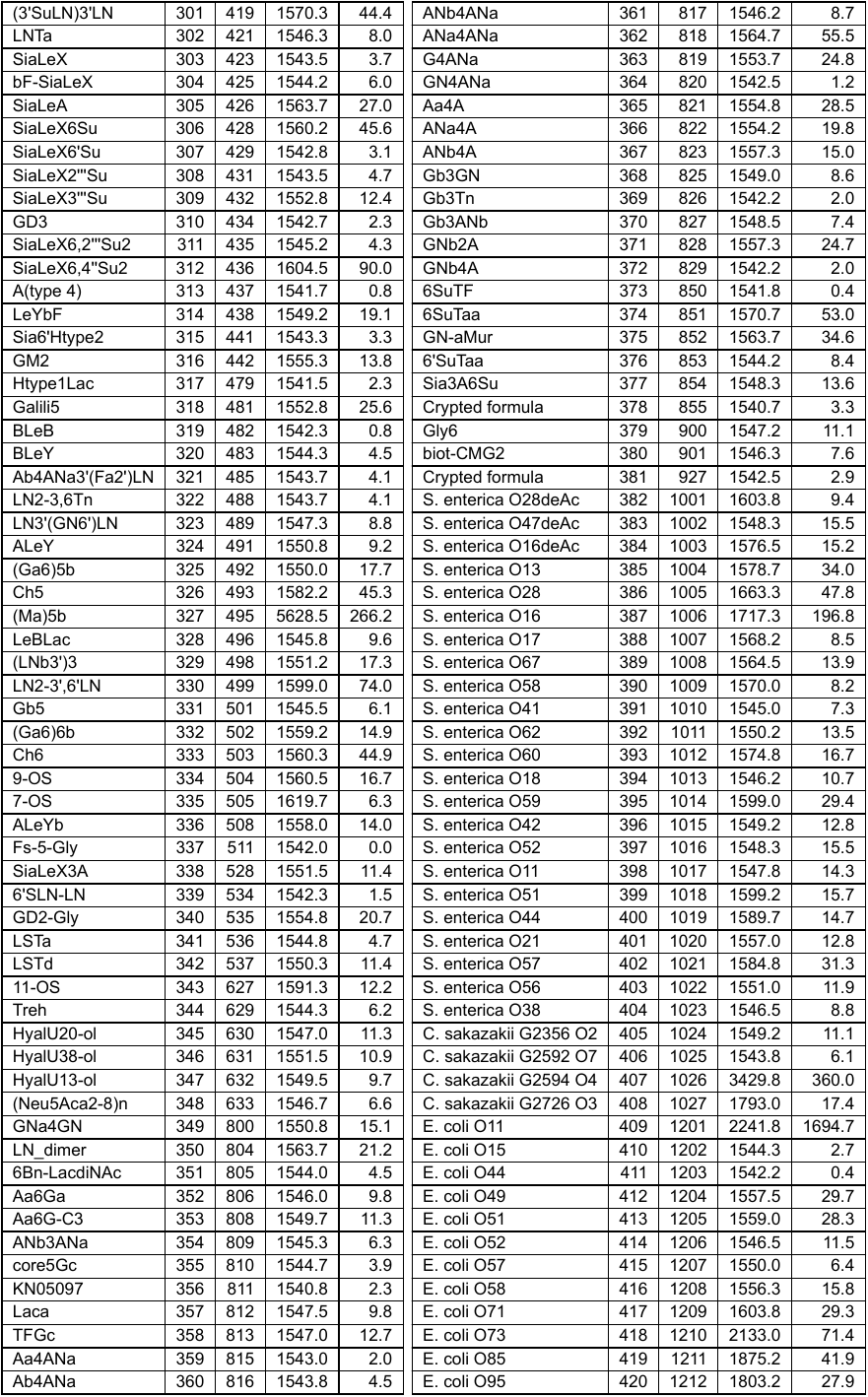

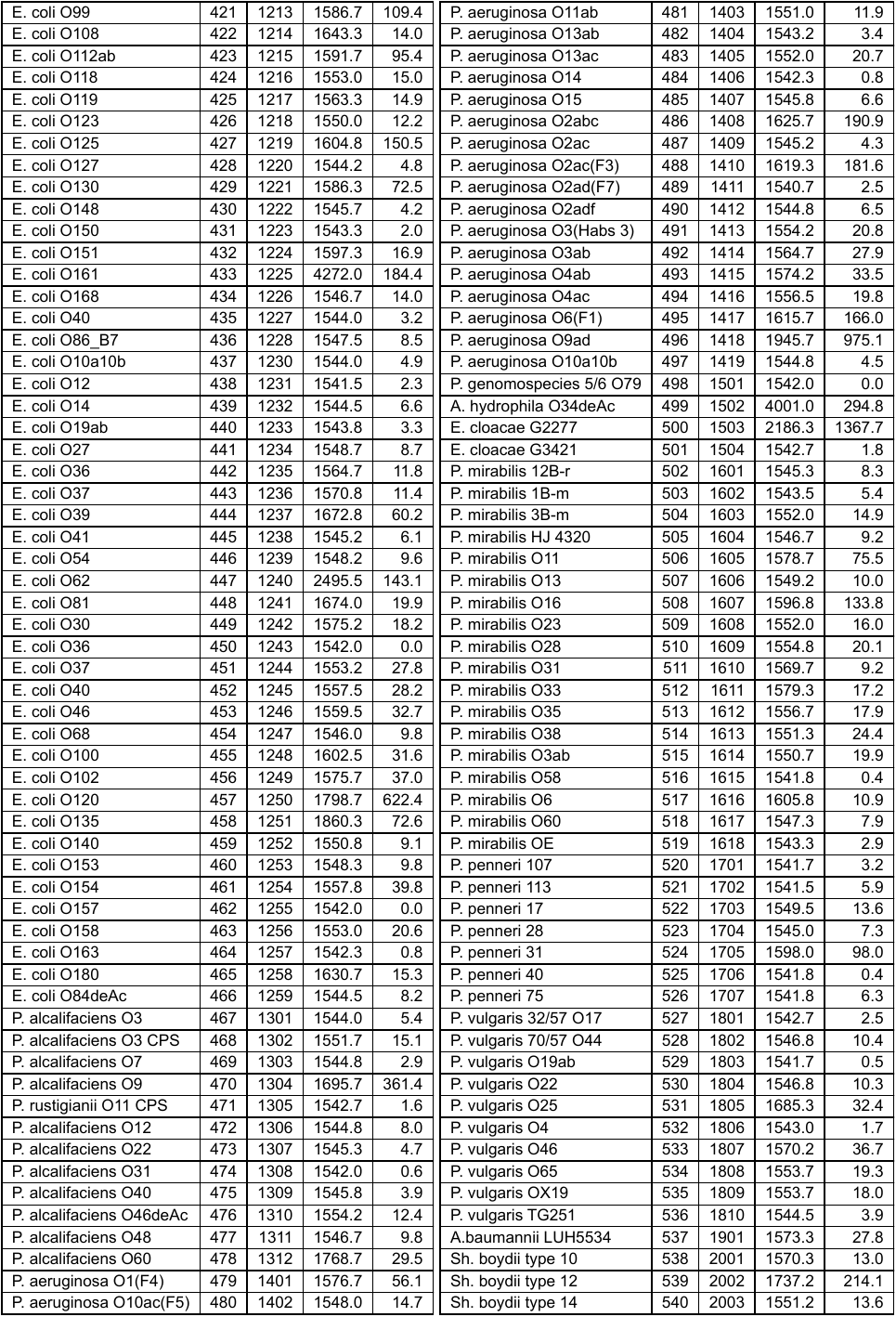

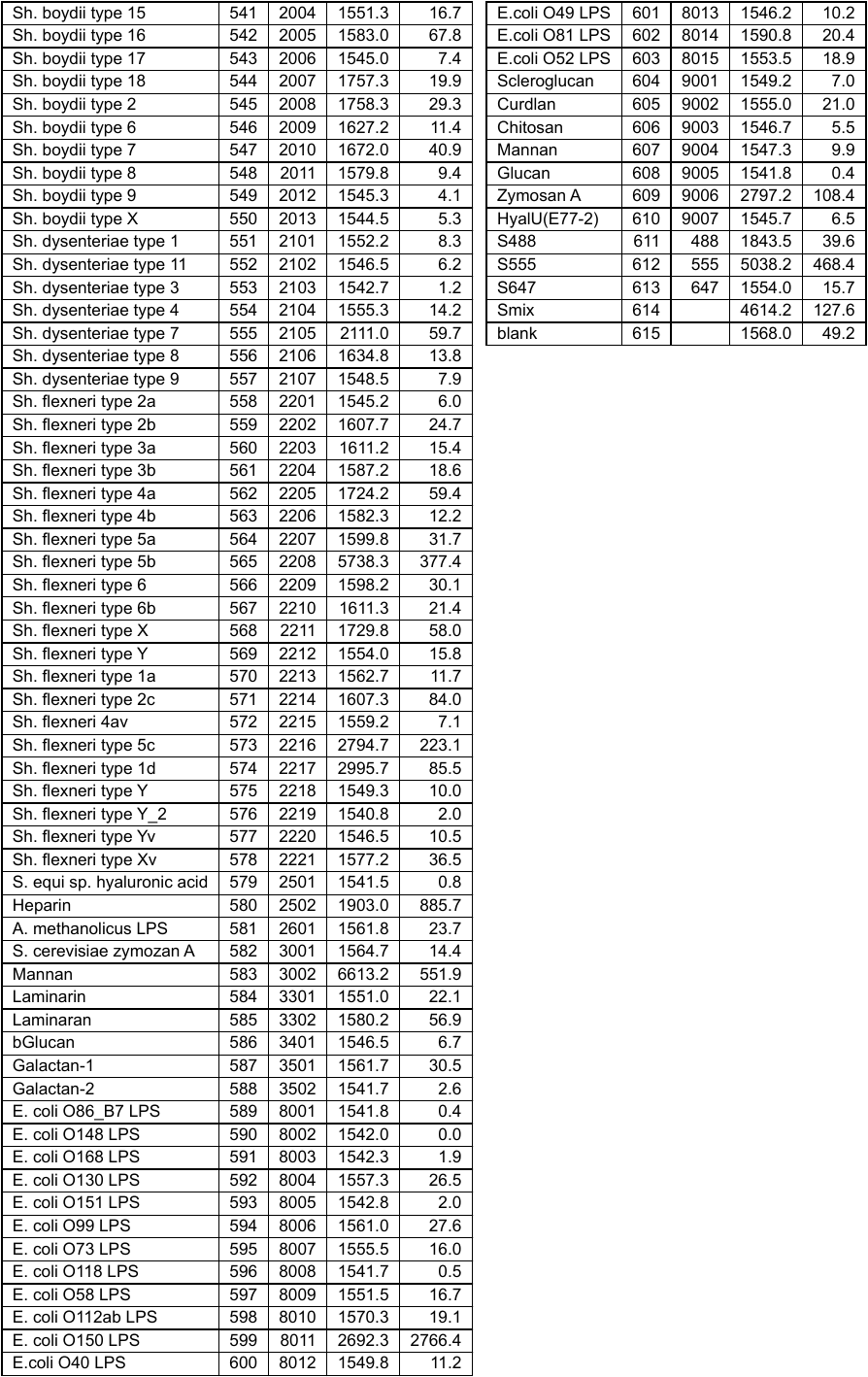

**Trypsin digest**

1 µg of S-protein (6.67 fmol) was dissolved in 50 mM aqueous (NH_4_)_2_CO_3_ pH 8. Disulfide bonds were reduced by adding 1 µL of 45 mM dithiothreitol (DTT) solution and incubation at 50 °C for 30 min. Cysteines were alkylated by adding 1 µL of 100 mM iodoacetamide solution and incubation for 15 min at r.t. in the dark. Excess iodoacetamide was destroyed by adding 2 µL 45 mM DTT solution and incubation for 30 min at 50 °C. 20 ng trypsin sequencing grade (TRYPSEQM-RO, Roche) were added and the mixture was incubated for 24 h at 37 °C. Formic acid to a final concentration of 20% v/v was added before sample preparation was done.

**Sample preparation**

Peptides from the trypsin digest were concentrated and desalted with ZipTips (Millipore® ZipTips C18). After washing the ZipTip with acetonitrile and water by pipetting, the peptides have been retained on the ZipTip by pipetting (25 steps) from the solution with the digested S-protein. The ZipTip was washed with 0.5% v/v formic acid solution at least 10 times to perform the desalting and washing of the peptides. Peptides were then eluted with acetonitrile/ 0.5% v/v formic acid solution 1:1 v/v first, and in the second elution step with acetonitrile. The final elution fraction had a volume of 16 µL. 0.5 µL of the elution fraction was spotted on a MTP AnchorChip 384 BC target (Bruker Daltonics, Germany) and dried under vacuum. 1 µL of α-cyano-4-hydroxycinnamic acid (Bruker Daltonics, Germany) dissolved in acetonitrile/0.5% formic acid 1:1 v/v to a concentration of 1mg/mL was spotted onto the dried droplet of peptide elution fraction. The α-cyano-4-hydroxycinnamic acid solution droplet was dried under vacuum.

**MALDI-MS method**

MALDI-MS was performed on a Bruker UltrafleXtreme. The measurement was performed in positive reflector mode with a *m/z* range from 400- 3500 Da. Ion source parameters were: 19.97 kV for Ion source 1; 17.97 kV for Ion source 2; 7.25 kV for the lens. Reflector parameters were: 21.33 kV for reflector 1, 10.68 kV for reflector 2.

**Spectra and peaklist processing**

Processing was done with Compass for flexSeries 1.4, flexAnalysis Version 3.4, Bruker BioTools 3.2 SR7 incl. digest calculating software (Bruker Daltonics, Germany).

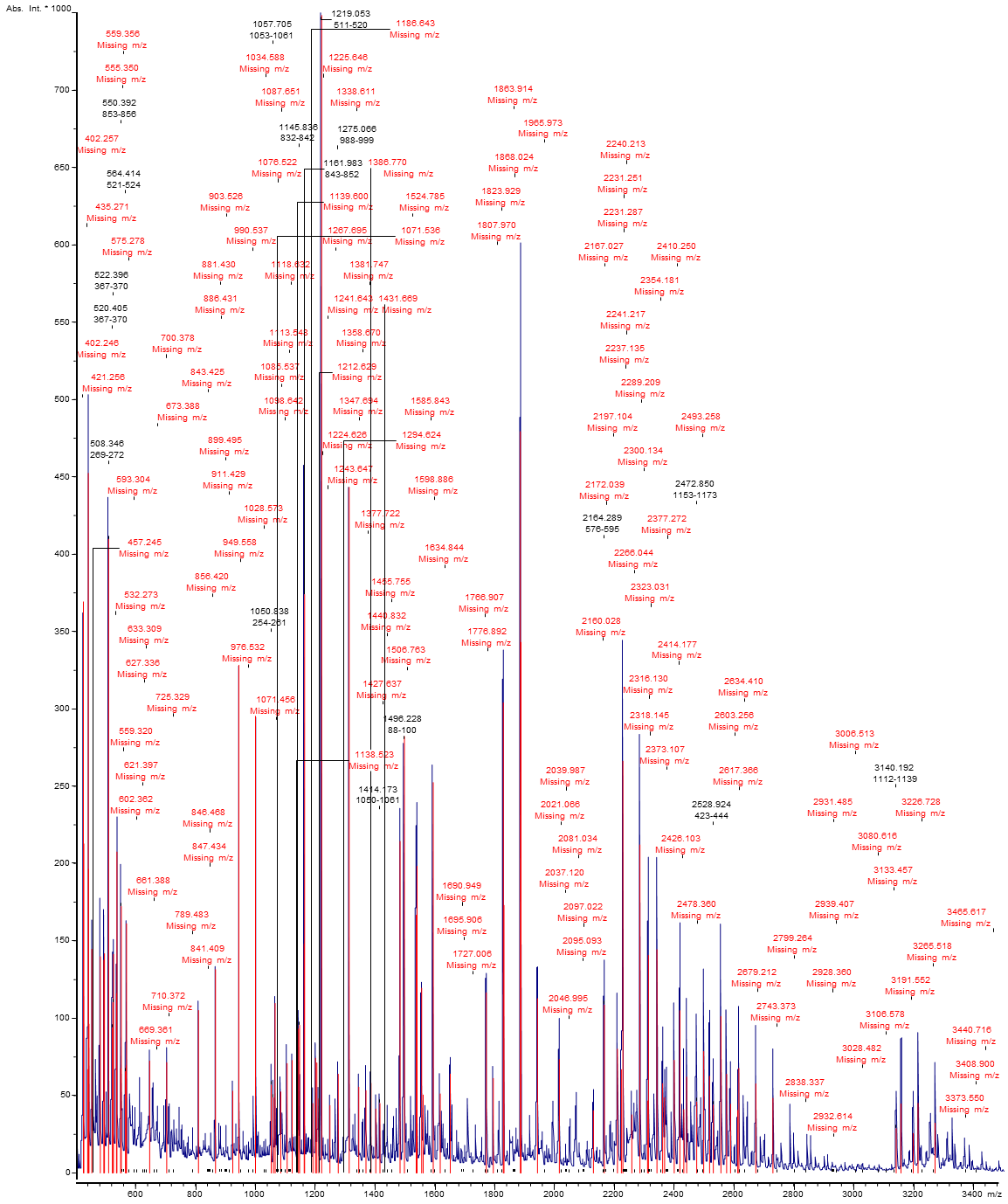

**Figure S5.** Mass spectrum of S-protein after trypsin digest.

**Table S3.** Parameter input into post processing software with coverage output results.

| Sequence name | S-protein |
| --- | --- |
| MH+ (mono) | 1.008 |
| Tolerance | 0.5 Da |
| Number of Peaks | 100 |
| Smoothing algorithm; parameters | SavitzkyGolay; width> 0.2 mz, 1 cycle |
| Intensity Coverage | 20.8% (2702849 cnts) |
| Sequence Coverage MS | 14.3% |

**Table S4.** Mass list, 100 peaks (most intense, S/N ≥ 6) with corresponding intensity.

| **Peak** | **Mass** | **Intensity** | **Peak** | **Mass** | **Intensity** | **Peak** | **Mass** | **Intensity** |
| --- | --- | --- | --- | --- | --- | --- | --- | --- |
| 1 | 424.198 | 369342.106 | 2 | 426.178 | 212881.878 | 3 | 438.201 | 66987.220 |
| 4 | 440.170 | 452432.421 | 5 | 440.910 | 196452.585 | 6 | 442.934 | 96438.576 |
| 7 | 452.260 | 144817.073 | 8 | 480.291 | 139939.326 | 9 | 492.343 | 137679.257 |
| 10 | 494.327 | 141897.901 | 11 | 508.346 | 409873.323 | 12 | 520.405 | 93739.228 |
| 13 | 522.396 | 110235.355 | 14 | 524.140 | 141505.786 | 15 | 534.388 | 116292.715 |
| 16 | 536.401 | 207728.114 | 17 | 548.400 | 67641.771 | 18 | 550.392 | 172478.265 |
| 19 | 564.414 | 65701.219 | 20 | 566.396 | 69595.314 | 21 | 568.123 | 161263.672 |
| 22 | 645.338 | 72564.830 | 23 | 702.388 | 71508.695 | 24 | 808.474 | 105219.826 |
| 25 | 865.520 | 131662.373 | 26 | 922.578 | 53055.497 | 27 | 943.609 | 328293.682 |
| 28 | 1000.662 | 297875.365 | 29 | 1050.838 | 56968.981 | 30 | 1057.705 | 48796.229 |
| 31 | 1063.755 | 109891.422 | 32 | 1082.716 | 52592.986 | 33 | 1103.781 | 71005.578 |
| 34 | 1120.787 | 72998.400 | 35 | 1141.773 | 93313.842 | 36 | 1145.836 | 95703.482 |
| 37 | 1160.872 | 148294.343 | 38 | 1161.983 | 374406.931 | 39 | 1189.810 | 45114.097 |
| 40 | 1198.842 | 74333.618 | 41 | 1202.917 | 71457.559 | 42 | 1209.898 | 48214.681 |
| 43 | 1219.053 | 748987.459 | 44 | 1246.882 | 43832.178 | 45 | 1275.066 | 64044.244 |
| 46 | 1311.100 | 443861.873 | 47 | 1343.110 | 55736.719 | 48 | 1367.110 | 53449.342 |
| 49 | 1384.174 | 58301.349 | 50 | 1400.155 | 41655.281 | 51 | 1414.173 | 45179.887 |
| 52 | 1481.405 | 214670.669 | 53 | 1496.228 | 282504.724 | 54 | 1535.314 | 166900.540 |
| 55 | 1538.480 | 198560.090 | 56 | 1553.316 | 119726.828 | 57 | 1558.323 | 50664.299 |
| 58 | 1592.386 | 252766.145 | 59 | 1615.426 | 51318.716 | 60 | 1649.424 | 64230.213 |
| 61 | 1769.639 | 116735.219 | 62 | 1792.726 | 61428.608 | 63 | 1826.715 | 304378.948 |
| 64 | 1828.858 | 173256.830 | 65 | 1883.795 | 479819.617 | 66 | 1885.949 | 343481.359 |
| 67 | 1940.908 | 113007.621 | 68 | 2014.193 | 73599.304 | 69 | 2128.329 | 40535.149 |
| 70 | 2164.289 | 108932.055 | 71 | 2208.299 | 80319.886 | 72 | 2221.385 | 67068.837 |
| 73 | 2226.442 | 266592.456 | 74 | 2256.457 | 37860.791 | 75 | 2283.532 | 212421.297 |
| 76 | 2308.656 | 46901.108 | 77 | 2311.566 | 141026.901 | 78 | 2340.628 | 144676.714 |
| 79 | 2359.669 | 57983.426 | 80 | 2365.621 | 49020.066 | 81 | 2397.719 | 73095.576 |
| 82 | 2416.779 | 105238.858 | 83 | 2429.797 | 41138.356 | 84 | 2439.841 | 62292.896 |
| 85 | 2472.850 | 46140.401 | 86 | 2496.902 | 78851.267 | 87 | 2515.986 | 62771.486 |
| 88 | 2528.924 | 44048.067 | 89 | 2554.031 | 101355.494 | 90 | 2573.109 | 64037.837 |
| 91 | 2586.050 | 44722.672 | 92 | 2610.116 | 40815.796 | 93 | 2614.074 | 67888.660 |
| 94 | 2671.169 | 57877.876 | 95 | 2728.260 | 48472.611 | 96 | 3140.192 | 28848.047 |
| 97 | 3156.203 | 44899.251 | 98 | 3197.270 | 25743.499 | 99 | 3213.333 | 45289.074 |
| 100 | 3270.487 | 32470.117 |  |  |  |  |  |  |

**Primary sequence of translated S-protein from expression vector, including signal sequence and His-tag (italic). The sequence of the viral protein is depicted in bold face**:

MGILPSPGMPALLSLVSLLSVLLMGCVAEGSWSHPQFEKSG*HHHHHHHH*DYDIPSSLEVLFQGPGS**MFVFLVLLPLVSSQCVNLTTRTQLPPAYTNSFTRGVYYPDKVFRSSVLHSTQDLFLPFFSNVTWFHAIHVSGTNGTKRFDNPVLPFNDGVYFASTEKSNIIRGWIFGTTLDSKTQSLLIVNNATNVVIKVCEFQFCNDPFLGVYYHKNNKSWMESEFRVYSSANNCTFEYVSQPFLMDLEGKQGNFKNLREFVFKNIDGYFKIYSKHTPINLVRDLPQGFSALEPLVDLPIGINITRFQTLLALHRSYLTPGDSSSGWTAGAAAYYVGYLQPRTFLLKYNENGTITDAVDCALDPLSETKCTLKSFTVEKGIYQTSNFRVQPTESIVRFPNITNLCPFGEVFNATRFASVYAWNRKRISNCVADYSVLYNSASFSTFKCYGVSPTKLNDLCFTNVYADSFVIRGDEVRQIAPGQTGKIADYNYKLPDDFTGCVIAWNSNNLDSKVGGNYNYLYRLFRKSNLKPFERDISTEIYQAGSTPCNGVEGFNCYFPLQSYGFQPTNGVGYQPYRVVVLSFELLHAPATVCGPKKSTNLVKNKCVNFNFNGLTGTGVLTESNKKFLPFQQFGRDIADTTDAVRDPQTLEILDITPCSFGGVSVITPGTNTSNQVAVLYQDVNCTEVPVAIHADQLTPTWRVYSTGSNVFQTRAGCLIGAEHVNNSYECDIPIGAGICASYQTQTNSPRRARSVASQSIIAYTMSLGAENSVAYSNNSIAIPTNFTISVTTEILPVSMTKTSVDCTMYICGDSTECSNLLLQYGSFCTQLNRALTGIAVEQDKNTQEVFAQVKQIYKTPPIKDFGGFNFSQILPDPSKPSKRSFIEDLLFNKVTLADAGFIKQYGDCLGDIAARDLICAQKFNGLTVLPPLLTDEMIAQYTSALLAGTITSGWTFGAGAALQIPFAMQMAYRFNGIGVTQNVLYENQKLIANQFNSAIGKIQDSLSSTASALGKLQDVVNQNAQALNTLVKQLSSNFGAISSVLNDILSRLDKVEAEVQIDRLITGRLQSLQTYVTQQLIRAAEIRASANLAATKMSECVLGQSKRVDFCGKGYHLMSFPQSAPHGVVFLHVTYVPAQEKNFTTAPAICHDGKAHFPREGVFVSNGTHWFVTQRNFYEPQIITTDNTFVSGNCDVVIGIVNNTVYDPLQPELDSFKEELDKYFKNHTSPDVDLGDISGINASVVNIQKEIDRLNEVAKNLNESLIDLQELGKYEQYIKW**PLER
